# Creating and leveraging bespoke large-scale knowledge graphs for comparative genomics and multi-omics drug discovery with SocialGene

**DOI:** 10.1101/2024.08.16.608329

**Authors:** Chase M. Clark, Jason C. Kwan

## Abstract

The rapid expansion of multi-omics data has transformed biological research, offering unprecedented opportunities to explore complex genomic relationships across diverse organisms. However, the vast volume and heterogeneity of these datasets presents significant challenges for analyses. Here we introduce SocialGene, a comprehensive software suite designed to collect, analyze, and organize multi-omics data into structured knowledge graphs, with the ability to handle small projects to repository-scale analyses. Originally developed to enhance genome mining for natural product drug discovery, SocialGene has been effective across various applications, including functional genomics, evolutionary studies, and systems biology. SocialGene’s concerted Python and Nextflow libraries streamline data ingestion, manipulation, aggregation, and analysis, culminating in a custom Neo4j database. The software not only facilitates the exploration of genomic synteny but also provides a foundational knowledge graph supporting the integration of additional diverse datasets and the development of advanced search engines and analyses. This manuscript introduces some of SocialGene’s capabilities through brief case studies including targeted genome mining for drug discovery, accelerated searches for similar and distantly related biosynthetic gene clusters in biobank-available organisms, integration of chemical and analytical data, and more. SocialGene is free, open-source, MIT-licensed, designed for adaptability and extension, and available from <u>github.com/socialgene</u>.

## Introduction

The advent of large-scale multi-omics datasets has ushered in a new era in biological research. However, the volume and complexity of datasets present significant challenges for their analysis. Here we present SocialGene, a suite of software for computing and organizing multi-omics data—including genomics, metabolomics, and more—into structured knowledge graphs^1^ ranging from small to repository-scale. While SocialGene is versatile and applicable across a broad range of disciplines, its initial development was motivated by the need to enhance genome mining for natural product drug discovery, the primary focus of this introductory manuscript.

Searching genomes for proteins of similar function and synteny (defined hereafter as a set of collinear, putatively-orthologous genes) is an essential task across multiple scientific disciplines. In natural product drug discovery, the focus centers on biosynthetic gene clusters (BGCs) that encode for the biosynthesis of specialized metabolites (SM). Developing new methods for identifying and targeting orthologous BGCs across vast sets of public and private genomes can enable a number of applications including finding additional sources of SM and their chemical analogs.

This is especially important, and challenging, for BGCs that encode for the biosynthesis of medicinally-important SM that have so far only been observed within the metagenome-assembled genomes (MAGs) of microbial obligate symbionts.^2,3^ Obligate symbionts often have reduced genomes^4,5^ and are recalcitrant to isolation and cultivation, necessitating the study of their SMs through chemical extraction of the holobiont or genetic engineering of predicted BGCs into heterologous hosts. These processes are resource-intensive but often necessary due to the difficulty and economics of chemically synthesizing many SMs. Several examples of the resources required to obtain SM from microbial endosymbionts include the antifungal SM lagriamide (symbiont producer– *Burkholderia gladioli*^6^), which required collecting 28,000 *Lagria villosa* beetle eggs to recover 600 µg of compound.^6^ The anticancer (and beetle-defense) compound pederin (symbiont producer– *Pseudomonas* sp.^7^) whose structure determination required the field collection of 25 million beetles (100 kg) over seven years, resulting in "many hospitalizations".^8^ The anticancer SM bryostatin (symbiont *Candidatus* Endobugula sertula^9^) which required 13,000 kg of *Bugula neritina* to produce 18 g of bryostatin 1 for clinical trials.^10,11^ The anticancer compound halichondrin B (proposed to be symbiont produced), which required 600 kg of the marine sponge *Halichondria okadai* for structure elucidation^12^ and an additional 1,000 kg collection of *Lissodendoryx* to recover approximately 300 mg of pure halichondrin for clinical trials.^11^ And the anticancer compound ET743 (symbiont producer–*Candidatus* Endoecteinascidia frumentensis^13^), which required aquaculture of 100,000 kg of the tunicate *Ecteinascidia turbinata* to recover the 100 g of compound needed up to Phase 2 of clinical trials.^14^ When the BGC of a metagenomic SMs is known or suspected, finding similar BGCs in cultured strains could be economically advantageous and influence the decision or speed of pursuing clinical trials. Given the non-generalizable and lengthy process of finding suitable vectors and hosts for genetic recombination, we sought a scalable framework for searching for related proteins and BGCs across previously cultured organisms, independent of BGC prediction frameworks, and where sequence identity and synteny might be low due to evolutionary distance.

Existing tools for finding similar BGCs (e.g. clusterblast^15^, MultiGeneBlast^16^, cblaster^17^, FlaGs^18^, CAGECAT^19^, BiG-SLiCE^20^ etc.) have proven valuable but had limitations for our use cases. Some require predicting BGCs within the target genomes and restricting searches to those BGC regions. However, current ensemble and machine learning BGC predictors (e.g. antiSMASH^15^, deepBGC^21^, GECCO^22^, etc.; review by Kim et al^23^) have a limited ability to detect BGCs not represented in their rule sets or training data, or when split across a sequence(s), necessitating a full-genome search. Others don’t scale well or rely on BLAST^24^ sequence similarity, which has limited ability to find distant homologs, particularly when search results are restrained and there are many homologs in the target database. As a result, we needed the ability to compare hundreds of millions of proteins and their positions across hundreds of thousands of genomes, in near real-time, while retaining the ability to discover low sequence identity homologs.

Protein similarity is often determined through sequence-sequence alignment tools (e.g. BLAST^24^, DIAMOND^25,26^, MMseqs2^27^, etc.) which, though incredibly fast, have a limit of detection for low sequence identity homologs, run considerably slower in high sensitivity modes, and often produce a burdensome number of matches when searching large target databases. This has forced a limitation in many current BGC search tools to only consider the top-n results of searches. Another approach is sequence-model alignment (e.g. HMMER^28^, HH-suite^29^, etc.) which provides detection of low sequence identity homologs but is too slow and compute-intensive for just-in-time annotation at repository scale. As both approaches are often needed, we aimed to develop a method that leverages each but performs the majority of the computation upfront, conducting dynamic searches over the stored results.

To that end, we created SocialGene, a suite of software projects centered around three elements: 1) a Python^30^ library that defines and controls the majority of data transformations and database interactions 2) a Nextflow^31^ workflow, written using nf-core^32^ templating and standards, that allows users to reproducibly create Neo4j graph databases using input proteins and/or genomes and 3) a Neo4j graph database, created in the final step of the Nextflow workflow, that stores and organizes the data, facilitating complex queries and analyses. The Nextflow workflow and Python library enable annotating proteins using profile hidden Markov models (pHMMs), clustering proteins with MMseqs2, and/or creating all-vs-all protein similarity networks with DIAMOND BLASTp. An option is available to run antiSMASH v7^33^ on all input genomes, allowing predicted BGC regions to be extracted and incorporated into the database. Additionally, the software and database schema were written modularly to allow users with programming experience the ability to extend the graph and perform custom analyses.

In this manuscript we present the software for the first time, as well as a limited set of potential use cases. This includes searching thousands of known BGCs against more than 343,000 RefSeq^34^ genomes, targeted genome mining for protein domains and functional co-occurrence, developing query strategies for drug discovery, linking Minimum Information about a Biosynthetic Gene cluster (MIBiG)^35^ BGCs to chemicals in NPAtlas^36^, linking genomes to LC-MS/MS features, Global Natural Products Social Molecular Networking (GNPS)^37^ clusters, and reference libraries.

As a resource for the community, we have precomputed SocialGene databases of various sizes, based on RefSeq as of November 14, 2023. This includes a SocialGene database computed over all RefSeq genomes, specific subsets of Actinobacteria, *Streptomyces* spp., and *Micromonospora* spp. genomes, and a database of 2,103,244 antiSMASH 7.0^33^ predicted BGCs– one of the largest public BGC compendia to date. Additionally, we provide an online and interactive BGC atlas. The atlas contains the results of using SocialGene to search the full size SocialGene RefSeq database for similar BGCS to each of the 2,502 BGCs in MIBiG^35^, but restricted to the >27,000 genomes associated with a strain available from a culture collection.

## Methods

### Nextflow workflow

Nextflow^31^ is a domain-specific language for producing reproducible scientific workflows. Nextflow was chosen for the promise of creating a single SocialGene workflow that would provide reproducibility, parallelism, checkpointing, and ability to run on local and cloud computing platforms. To provide standardization, SocialGene’s database-building workflow was designed to nf-core^32^ standards. nf-core is a “framework for community-curated bioinformatics pipelines”^32^ and, while SocialGene was not submitted as an official nf-core workflow, it was built using the framework and therefore benefits from the surrounding tooling. This includes a GUI for launching the workflow and the ability to interface with nf-core "tools", "modules", "subworkflows", etc. Additionally, the workflow can be run using Seqera’s Nextflow Tower, an online Nextflow workflow orchestrator. SocialGene’s Nextflow workflow (github.com/socialgene/sgnf) handles downloading data from a number of sources (e.g. NCBI genomes, MIBiG BGCs, multiple public pHMM databases, etc.), the extraction, transformation, and loading (ETL) of input and computed data, and culminates in the building of a custom Neo4j graph database (Fig. 1 and Supplementary Fig. 1). The SocialGene Nextflow workflow and Python library were designed modularly so that users can choose to run any or all analyses. The configuration files used for creating the Neo4j databases in this manuscript are available within the archived codebase. As of writing, SocialGene was built against Nextflow version 24; and nf-core tools template version 2.10.

**Figure 1.**
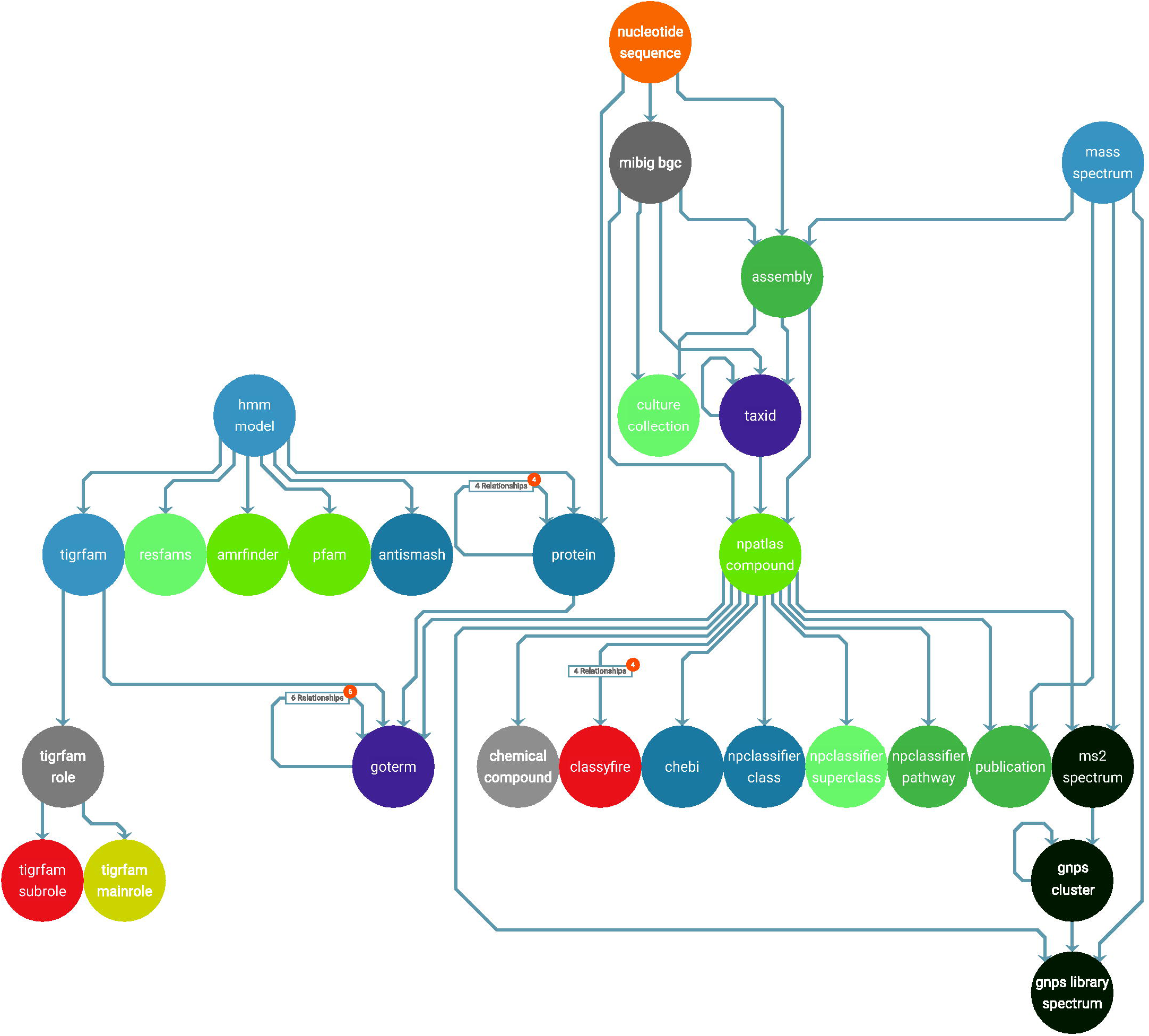
A partial schema illustration of a SocialGene database, showing nodes (circles) and their relationships (lines between circles). The visualization was auto-generated and formatted by connecting the RefSeq-based Neo4j database to yFiles’^39^ Neo4j Explorer. A high-resolution version is available (see Data and Code Availability), and an up-to-date, auto-generated, full node-relationship schema is available in SocialGene’s online documentation.

### Python library

The Nextflow workflow contains several independent Python scripts but also makes use of command line entry points defined within the SocialGene python library (github.com/socialgene/sgpy; pypi.org/project/socialgene). The library was written with entry points for limited use as a command line tool, as utilized in the Nextflow workflow, and as a Python library directly for a number of bio- and cheminformatic tasks. All code changes are checked for breaking changes (pytest) and code style (Flake8 and Black) through continuous integration and continuous delivery (CI/CD) workflows using GitHub Actions and "Release Please"^38^ which automate new releases and deployments to PyPI. Test coverage is monitored with Codecov. With the SocialGene software split across several git repositories and software languages it was important to coordinate a consistent set of parameters when using each (e.g. parameters passed to HMMER’s hmmsearch when creating a SocialGene database and when annotating a query protein later). To maintain consistent settings across database creation (via Nextflow), notebook analysis (Python), and future interfaces (Django), the Python library contains a file of environment variables “common_parameters.env” which are read and modified at runtime from within Nextflow, Django, etc. These parameters are also saved within the Neo4j database at the time of creation.

### Neo4j graph database

Neo4j is a company that maintains graph database software of the same name. Neo4j maintains support of docker images of community and enterprise editions, drivers in popular languages, and in-database graph data science and machine learning libraries. The SocialGene Nextflow workflow and Python library automate the creation of bespoke Neo4j databases, which can then be interrogated directly in a web browser, with the SocialGene Python library, Cytoscape, or other third party tools. While SocialGene makes use of Neo4j graph databases, the Nextflow workflow gathers all intermediate files as tab-separated flat files that could be imported into an alternate database system.

### Input genomes

The Nextflow workflow can download genomes from NCBI or use local GenBank files and/or protein FASTA files. To identify redundant proteins, as well as provide a consistent, cross-source sequence-based identifier, SocialGene uses sequence hashes as universal identifiers. When genomes are provided in GenBank format SocialGene uses BioPython^40^ and custom scripts to parse genome and sequence data. Additionally, we have found highly-relevant pseudogenes within BGCs (and elsewhere) which lack translated sequence data in GenBank files. With this observation, and recent studies showing some portion of PGAP-labeled^41^ pseudogenes are misassembled coding genes,^42^ we decided to attempt to include annotated pseudogenes. Therefore, SocialGene attempts to include pseudogene content via extracting the relevant nucleotide sequence and employing BioPython’s Bio.Seq.translate. As some pseudogenes simply contain a potential early stop codon(s) these may also be physiologically relevant via translational read-through^43^ and other mechanisms. However, the correct translation of pseudogenes that aren’t transcribed or the incorrect translation of pseudogenes is a data inclusion bias users should be aware of. SocialGene tracks which sequences were derived from pseudogenes by prepending "pseudo_" to the locus description, which can be used to filter results in the SocialGene database. Additionally, if available in the GenBank file, SocialGene will attempt to include the reason the gene was marked as pseudo, (e.g internal stop, frameshift, etc.).

### Representing proteins as hashes

SocialGene Neo4j database protein entries use sha512t24u^44^ hashes as universal identifiers but are also assigned a CRC64 hash for fast cross-referencing with UniProt.^44,45^ Hashing is a process that takes an input string of characters and transforms it into a uniquely identifiable hash. This is often used to assign a short, unique identifier to a large quantity of information. For example, the human protein titin (UniprotKB^46^ Q8WZ42) contains 34,350 amino acids but can be represented by its CRC64 hash: DEB216410AD560D9. Different hashing algorithms have different probabilities for the scenario that two different inputs produce an identical hash (hash collision). While preparing this manuscript we switched to using CRC64 due to its use in UniProt which would provide the ability to crosslink, and link out to, UniProt information/resources^45^. However, it was discovered there were 1,704 hash collisions across UniParc (out of 517,621,195 total proteins) and, more concerning, tens of thousands of collisions when hashing SocialGene’s internally used string "{genbank_accession} {genbank_locus}" across all RefSeq nucleotide sequences. For this reason we switched back to using sha512t24u due to its predicted collision probability (no collisions were detected in UniParc or internal SocialGene identifiers), speed (∼2x SEGUID), and being url-safe.^44^

### Hidden Markov models and annotation

SocialGene’s Nextflow workflow can download and format pHMMs from any or all of antiSMASH^47^, AMRFinder^48^, BiG-SLiCE^20^, ClassiPhage^49^, Pfam^50^, PRISM^51^, Resfams^52^, and TIGRFAMs^53^; as well as user-provided pHMMs in HMMER3 format. The Python library reduces input pHMMs to a less-redundant set of models by hashing the models’ emissions and transitions and, for Pfam, uses only the latest version of a model. For example, if the user or combination of above reference databases try to include Pfam models PF00001.23 and PF00001.24, SocialGene will only annotate proteins with PF00001.24, and in the resulting Neo4j database will note that the PF00001.23 model was specified by the input source but PF00001.24 was used for domain prediction. For compatibility with HMMER’s hmmsearch, less-redundant models are output in two files, one for models with gathering cutoffs and one for models without. Within Nextflow, less-redundant fasta files are split into n-files and run against the two pHMM files in parallel using HMMER’s hmmsearch. Through extensive testing we found that, with fast hard drives, splitting an input fasta into multiple files, assigning 1 logical cpu to hmmsearch, and running in a highly parallel fashion provides the fastest results for this step.

### antiSMASH

SocialGene’s Nextflow workflow has the ability to annotate input genomes with antiSMASH version 7^33^. A custom Python script reduces resulting antiSMASH json files into a minimal JSONL file (newline-delimited JSON) that describes the assembly, locus, coordinates and minimal metadata for all predicted BGCs. While at first this may seem unnecessary, the gzipped tar archive of unmodified antiSMASH output for all successfully annotated RefSeq genomes was >1.5 TB. The gzipped, summarizing, gzipped minimal JSONL for the same was 86 MB, a >16,000x reduction in storage size.

### MMSeqs2

SocialGene’s Nextflow workflow performs cascaded clustering using MMseqs2. For example, clustering non-redundant proteins to 90% and 50% sequence identity first clusters proteins to 90%, followed by taking the 90% cluster representatives and clustering them to 50%. This is important because it means to find proteins in the database with less than 90%, but greater than 50%, sequence similarity will require a two-hop traversal, first traversing "MMSEQS_90" relationships then "MMSEQS_50". To allow users to cluster input proteins to multiple, custom identity levels the Nextflow module was written to take a delimited string of identity levels (e.g. ’90,70,50’, representing 90%, 70%, and 50% sequence identity). Depending on the number of proteins and clustering levels this process can require a significant amount of disk space and RAM (100s of GBs). This Nextflow process outputs a single flat file edge list representing the protein clusters, as well as MMseqs databases for each level.

### Hardware

Data was created/analyzed on either: "Desktop 1": single AMD® Ryzen 9 3900xt 12-core processor with 62 GB of RAM; or "Server 1": dual AMD® EPYC 7352 24-Core processors, with 1 TB RAM. Both machines used SABRENT 4 TB Rocket NVMe PCIe M.2 2280 as working drives. NCBI RefSeq genomes were stored on, and processed from, a Western Digital 18TB WDC_WD181KRYZ disk drive. Large scale pHMM annotations were computed using the University of Wisconsin-Madison’s Center for High Throughput Computing (CHTC) and the Open Science Grid (OSG). While not required, Neo4j database creation, initialization, and large-scale read/write benefit from fast hard drive storage.

### Scaling

Databases used in this manuscript were computed using a combination of Desktop 1, Server 1 and CHTC resources. For inputs over 1,000 genomes, data aggregation steps can be computed on a mid-tier laptop or desktop computer, but the non-distributed DIAMOND and MMseqs2 protein comparisons begin to require a high amount of RAM and it becomes best to shard the input FASTA and run the pHMM annotation step on a high throughput computing cluster, if available. Since Nextflow is not currently supported on the University of Wisconsin-Madison’s CHTC submit server, a flag "--htcondor" was created in SocialGene’s Nextflow workflow which signals for the organization and output of the bundled set of processed non-redundant fasta files, pHMMs, generated scripts, and instructions for submitting the jobs via HTCondor (but generalizable to other computing environments). The Nextflow workflow can then be run a second time with the "--htcondor" flag removed and the path to HMMER results provided to the command line flag "--domtblout_path". Adding the "--resume" flag allows this second run to continue where the first left off, reusing already completed computations. Utilizing this technique, combined with resources available through CHTC and OSG, has allowed us to create SocialGene databases with >340,000 genomes, requiring tens of thousands of CPU hours, in under two days, instead of months to years.

### Precomputed databases

To test SocialGene’s ability to scale to large collections of genomes, we ran the workflow on all genomes available in RefSeq (including non-bacterial). While it’s possible to use SocialGene to download all RefSeq genomes, doing so requires a substantial amount of disk space (>1.5 Tb) and thus we used an existing local copy of the 343,381 genomes, updated on November 14, 2023. The SocialGene Nextflow workflow was run on Server 1 with settings to annotate all genomes with antiSMASH 7^33^; annotate all non-redundant proteins with pHMMs from antiSMASH^33^, AMRFinder^48^, Pfam^50^, Resfams^52^, and TIGRFAM^53^; and cluster non-redundant proteins to 90%, 70%, 50%, 30% with MMSeqs2. To run hmmsearch on CHTC/OSG we instructed the Nextflow workflow to split the non-redundant protein FASTA into 3000 files (using SeqKit^54^ split). SocialGene’s Nextflow flag "--htcondor" then instructed the workflow to package the resulting FASTA files, two non-redundant pHMM model files (those with and without gathering cutoffs) and customized scripts, for submission with HTCondor. The resulting 6,000 hmmsearch jobs required 14,726 cpu hours to complete and the total workflow required approximately 17,000 CPU hours. This does not account for the more than 10,000 CPU hours to compute antiSMASH BGCs across all 343k genomes on Server 1. Apart from downloading input genomes and antiSMASH predictions, due to its parallel design, the workflow completed start-to-finish in less than 48 hours. However, it should be noted this is highly variable and dependent on the compute resources, especially the number of CPUs. Supplementary Table 1 shows the number of nodes and relationships in the resulting database. The full graph database occupies 650 GB of disk space and is available for download as a 220 GB Neo4j database dump (see Data and Code Availability). We are also making available a separate 30 GB SocialGene database of the more than two million antiSMASH predicted BGCs.

Three additional RefSeq databases (named "actinomycetota", "streptomyces", and "micromonospora") were created with the intention of providing smaller precomputed databases for those without access to adequate computational resources. Each was built independently with the SocialGene Nextflow workflow, making use of the NCBI Datasets module (e.g. --ncbi_datasets_command ’genome taxon "actinomycetota" --assembly-source refseq --exclude-atypical’). Databases in this manuscript have been labeled as version 2023_v0.4.1.

### Representing and linking chemistry

SocialGene has the ability to incorporate and crosslink non-redundant chemical compounds from a variety of sources, using RDKit^55^ and custom scripts. As of writing, redundancy is based on unique InChI^56^ strings, as most NP databases don’t contain more detailed structural information than InChI^56^ or SMILES^57^. Additionally, SocialGene links similar compounds within the database using an all-vs-all comparison of Morgan fingerprints^58^ (radius 2, 2048 bits) and Tanimoto similarity.

## Results and Discussion

### HMM outdegree accelerated BGC search

The SocialGene RefSeq database (version 2023_v0.4.1) contains data on >340 thousand genomes, >300 million non-redundant proteins, >25 thousand less-redundant pHMMs (from antiSMASH^47^, AMRFinder^48^, Pfam^50^, Resfams^52^, and TIGRFAMs^53^), and >840 million pHMM-to-protein annotations. Evenly distributed, this would result in 35,919 annotations per pHMM. But, as shown in Supplementary Fig. 2, the actual distribution of annotations per model is right-skewed and log-normal distributed, with a mean of 33,163 and median of 2,948.

SocialGene’s BGC search first annotates a query BGC’s proteins using the same pHMM models used to annotate proteins in the database, either pulling annotations from the database when the protein is present, or using HMMER^28^. To compare proteins by their pHMM annotations reduces the initial search space to the 25,566 pHMM nodes, but the number of outgoing relationships from pHMM nodes is over 847 million. Consequently, searches can quickly begin traversing an excessive percentage of the database. To alleviate this SocialGene’s BGC search algorithm first calculates, and sets as a node property, the outdegree of pHMM nodes. The input proteins are then prioritized by the lowest to highest summed outdegree of their pHMM annotation nodes.

The database is then searched for all proteins with similar domains (pHMM annotations). These similar proteins and their gene coordinates within all genomes are clustered and filtered in Python based on a threshold number of hits to the input BGC’s proteins. After filtering, the remaining nucleotide sequences are divided into multiple regions based on the user-specified ’break_bgc_on_gap_of’ parameter, which splits a nucleotide sequence where any region of the specified length has no hits to an input BGC protein. Regions are filtered again by a threshold number of hits to the input BGC proteins. Remaining regions are evaluated by reciprocal best hit (RBH) analysis using either DIAMOND BLASTp or pHMM annotation similarity (user-selected). The resulting putatively similar BGCs are then evaluated and ranked based on the similarity of RBH content (Jaccard) and order (Levenshtein) compared to the input BGC. The search can be done either within an interactive Python terminal or Jupyter notebook, enabling further computation, or as a standalone command line function which outputs a JSON file for visualization with clustermap.js^59^.

This outdegree prioritization can dramatically speed up a search and essentially prioritizes less common pHMM annotations (and, thereby, domains and proteins). However, this strategy can miss target clusters if the only related proteins between query and target BGCs are those excluded in the prioritized search. Fig. 2 provides a visual guide of this relationship of pHMMs (and their outdegree) to protein and nucleotide sequences (labeled as BGC in the figure). Further explanation is available in <u>Supplementary Text 1</u>.

**Figure 2.**
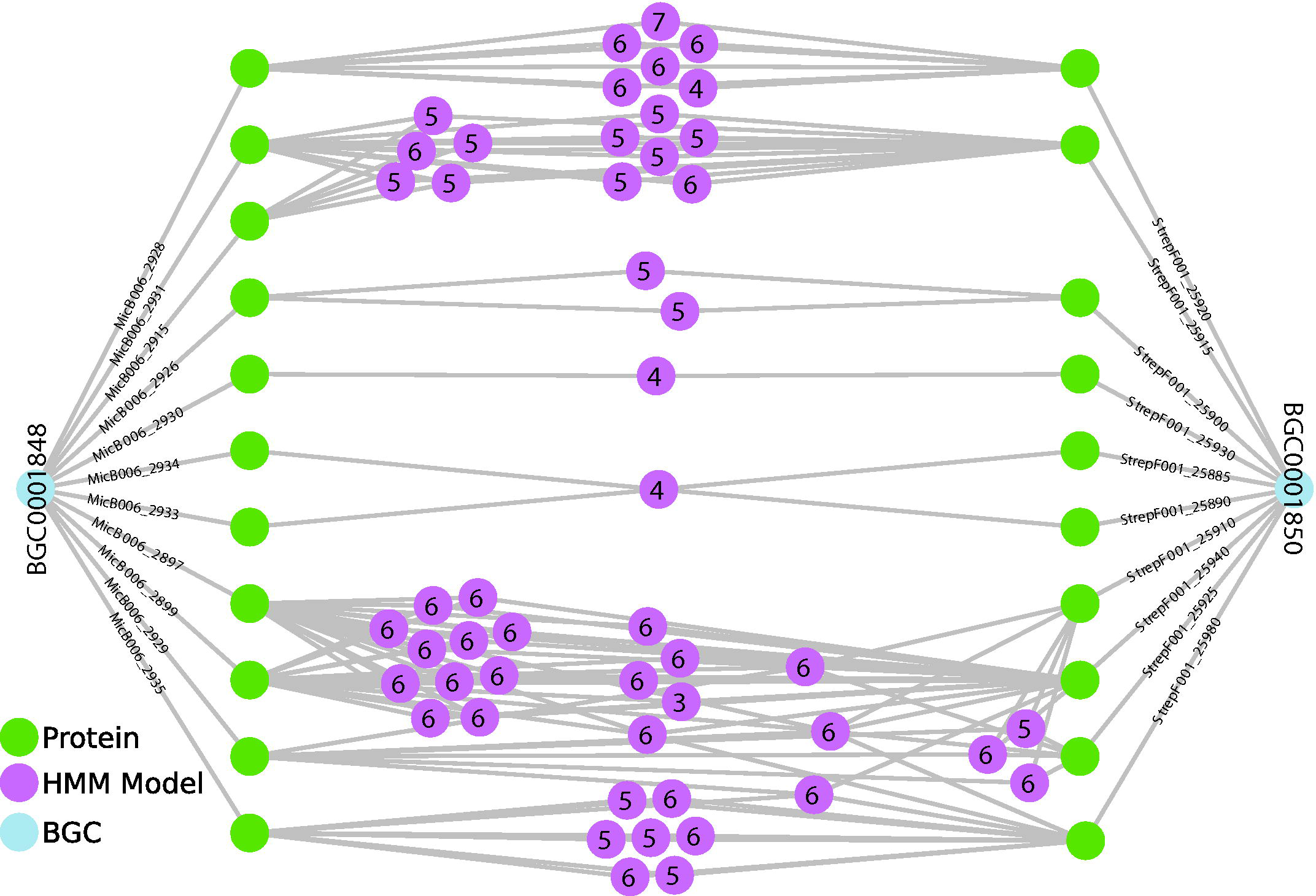
A simplified illustration of two BGCs from MIBiG, their encoded proteins, and shared pHMM annotations (gray lines). pHMMs models are labeled with numbers representing the log of their outdegree (e.g. 4 is approximately 1,000 relationships, 6 is approximately 100,000). Searching SocialGene databases for similar BGCs is accelerated by a first-pass search of annotations by low-outdegree pHMMs. The illustration also highlights the complexity of shared pHMM annotations between proteins within a single BGC. A comprehensive comparison of the two displayed BGCs was previously published by Braesel et. al.^60^

### Multiple methods are required for measuring protein similarity

To justify the protein similarity search strategy for biosynthetic gene clusters (BGCs) we explored the correlation between DIAMOND’s^25^ BLASTp protein-protein sequence identity scores, MMseqs2 clustering, and the Jaccard and Levenshtein similarity of HMMER^28^ pHMM annotations. While MMseqs2^27^ and DIAMOND were comparable (See Supplementary Figs. 3,4), there was little, if any, global correlation between pHMM annotations and DIAMOND BLASTp identities (Supplementary Fig. 5). This lack of correlation is due to the algorithms used for pHMM annotation similarity which don’t account for model or sequence coverage, or detailed domain position. For single domain proteins, perfectly-similar pHMM annotation often consists of only a single pHMM model annotation. Thus, while single domain proteins can have a range of sequence alignment identities they usually only have binary pHMM Jaccard and Levenshtein similarity scores.

For example, UniProtKB^46^ proteins Q8X5K5 and Q8XCP8 are encoded by *Escherichia coli* O157:H7 genes *lpfA* and *yfcQ*. Both proteins are potentially highly relevant to human health, as *lpfA* is part of the *lpfABCC’DE* fimbrial operon and has been shown to promote enterohemorrhagic *E. coli* cells’ interaction and adherence to eukaryotic cells.^61–66^ However, while *yfcQ* from the laboratory-cryptic *yfcOPQRSTUV* operon has been computationally inferred to also be a fimbrial-like adhesin protein, there have been limited studies on its role in pathogenesis or adhesion.^67–69^ While NCBI’s BLASTp^24,70^ was unable to align these protein sequences due to their low sequence similarity, their predicted AlphaFold^71^ 3D protein structures did align (see Supplementary Fig. 6). Additionally, when looking at pHMM annotations, over 80% of the AAs in both proteins were annotated by the PF00419.23 (Fimbrial) Pfam model. Therefore, a search strategy starting with one of these proteins would fail to find the other when using BLASTp, while a strategy employing pHMM annotations would succeed.

Conversely, a nearly-perfect BLASTp alignment doesn’t necessitate similar pHMM annotation. For example, UniProtKB^46^ A0A0H3JI96 and A0A0H3JGM8 are phage tail proteins encoded in the *Escherichia coli* O157:H7 genome. While BLASTp alignment revealed matches in 233 of 238 positions (97.9% identity), only a third of their pHMM annotations overlap in SocialGene’s RefSeq database (see <u>Supplementary Text 2</u>).

Therefore, it is important to consider using both sequence and model approaches to protein similarity in SocialGene. For large databases, we recommend utilizing SocialGene’s MMseqs2 cascaded clustering method rather than all-vs-all BLASTp, as the latter can result in an excessive number of relationships.

### Finding metagenomic, fragmented and multiple copy BGCs

Searching for metagenome-assembled genome (MAG) derived BGCs in the sequences of cultivated organisms is challenging for a variety of reasons including the sheer number of public genomes and the low quality of many MAG BGCs and public genomes. To examine the ability of SocialGene’s BGC search algorithm to look through hundreds of thousands of public genomes for metagenomic BGC homologs we looked at a recently verified example: lagriamide. Lagriamide is an antifungal SM whose BGC is encoded in the reduced genome of the *Lagria villosa* beetle endosymbiont, *Burkholderia gladioli* Lv-StB.^6,72^ Through a combination of individual BLASTp searches against NCBI’s nr database and manual bioinformatic analysis we and several laboratories recently collaborated to find two free-living strains of *Paraburkholderia acidicola* that contained a partial match to the metagenomic-derived BGC encoding for lagriamide.^6,72^

To evaluate SocialGene’s BGC search algorithm we took the MAG-derived lagriamide BGC (MIBiG BGC0001646) and ran the search against the SocialGene RefSeq database (343,381 public genomes) and were able to recover the aforementioned *P. acidicola* BGC (Supplementary Fig. 7).

SocialGene was also able to recover the BGC of the immunosuppressant SM rapamycin, even when fragmented and/or containing corrupted-genes, as shown in Supplementary Fig. 8. While SocialGene is able to find fragmented BGCs, the default search returns only the highest-scoring fragment due to limitations of plotting in clustermap.js^59^. Lastly, the BGC search function was able to recover the multiple integrations of a nybomycin-encoding plasmid in a genome engineered strain (Supplementary Fig. 9).

### Finding syntenic but distantly related BGCs

There are few examples of finding endosymbiont-derived, metagenomic BGCs in free-living relatives and few references of the extent of sequence divergence of orthologous BGCs over large evolutionary distances. While the lagriamide example above was reported to have 93.7% pairwise identity,^72^ the individual proteins in the public genome assembly have amino acid identities around 70 to 80%. And though we hypothesized the need to find syntenic BGCs where individual ortholog sequence similarities were low we were unsure if this existed in nature. Additionally, while finding collinear, putatively-orthologous genes is suggestive of common ancestry and conserved function it is important to consider the likelihood of convergent evolution, though the probability of the later assumedly decreases as the sequence similarity, count, and synteny of shared genes increases.

To test our hypothesis, SocialGene’s automated BGC search algorithm was used to search each of 2,502 MIBiG BGCs as queries against the entire SocialGene RefSeq database. While the algorithm uses pHMM annotations for the primary search, the lower bound of sequence identity was limited by the final step, where putative target BGCs were compared with reciprocal best hit (RBH) analysis using DIAMOND BLASTp in "ultra-sensitive" mode. Though confounded due to biases in the RefSeq and MIBiG databases,^73^ the majority of query BGCs that had targets with high synteny and low sequence similarity also had a large number of total hits (i.e. BGCs that are highly prevalent across RefSeq). These abundant BGC classes were similar to Cimermancic, Medema, Claesen et al’s observation of the widespread occurrence of "O-antigens, capsular polysaccharides, carotenoids and NRPS-independent siderophores".^74^

To that end we looked for the longest MIBiG BGC with the highest synteny and lowest median RBH identities. MIBiG BGC0000182 is a BGC from a *Pseudomonas fluorescens* bacterium with 36 protein-coding genes that encode the biosynthesis of the polyketide antibiotic pseudomonic acid A (mupirocin). Mupirocin is a clinically-important antibiotic that continues to be included on the World Health Organization’s List of Essential Medicines.^75^ Using SocialGene’s BGC search function, we searched the SocialGene RefSeq database for BGC0000182. While most of the resulting 17 target BGCs were highly similar, two had median RBH identity values of 73.8% and 58.5%, while still containing a RBH to every BGC0000182 protein (Fig. 3). While the strain with a median of 73.8% protein sequence identity was also a *Pseudomonas* sp., the strain with 58.5% median identity was *Chromobacterium* IIBBL 290-4, which belongs to a different taxonomic Class. The *Chromobacterium* sp. BGC was flanked by transposases (NKT35_RS10105/NKT35_RS10110 and NKT35_RS10295) suggesting potential mobility of the BGC. Interestingly, the region between transposons only contains 34 of the 36 proteins, with MupR and MupX homologs occurring directly adjacent to, but outside of, NKT35_RS10295 (a pseudo IS1380 family transposase). While *Pseudomonas* sp. QS1027 is a known producer of mupirocin.^76^ it is currently unknown whether *Chromobacterium* IIBBL 290-4 produces mupirocin or a mupirocin chemical analog.

**Figure 3.**
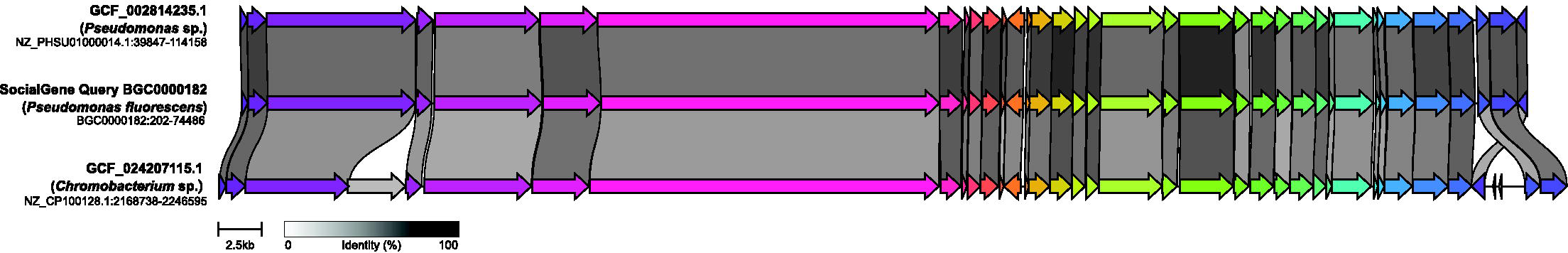
SocialGene’s BGC search function outputs a clustermap.js^59^ plot, as shown. The abridged plot here displays two target BGCs, obtained from searching >343,000 genomes for BGCs similar to the mupirocin-producing BGC (BGC0000182, middle row). Two result hits are displayed in the top and bottom rows, with median RBHs of 73.8% and 58.5% to BGC0000182, respectively, as calculated by DIAMOND BLASTp. Individual alignment identities are shown in Supplementary Fig. 10.

### Following the evolution of BGCs

While some BGCs have few matches (e.g. mupirocin, mentioned above), others are overrepresented due to organism bias in RefSeq, wider phylogenetic distribution, or both. Using the same search strategy as with mupirocin, but with BGC0000946 (*Vibrio parahaemolyticus* BGC encoding for vibrioferrin) as the query BGC, resulted in 6,577 complete and syntenic target BGCs across 6,571 genome assemblies, along with 4 MIBiG BGCs. These BGCs were distributed across 968 species, 81 genera, 46 families, 6 classes, and 3 phyla; with the median percent identities of RBHs occurring in a stepped gradient (Fig. 4).

**Figure 4.**
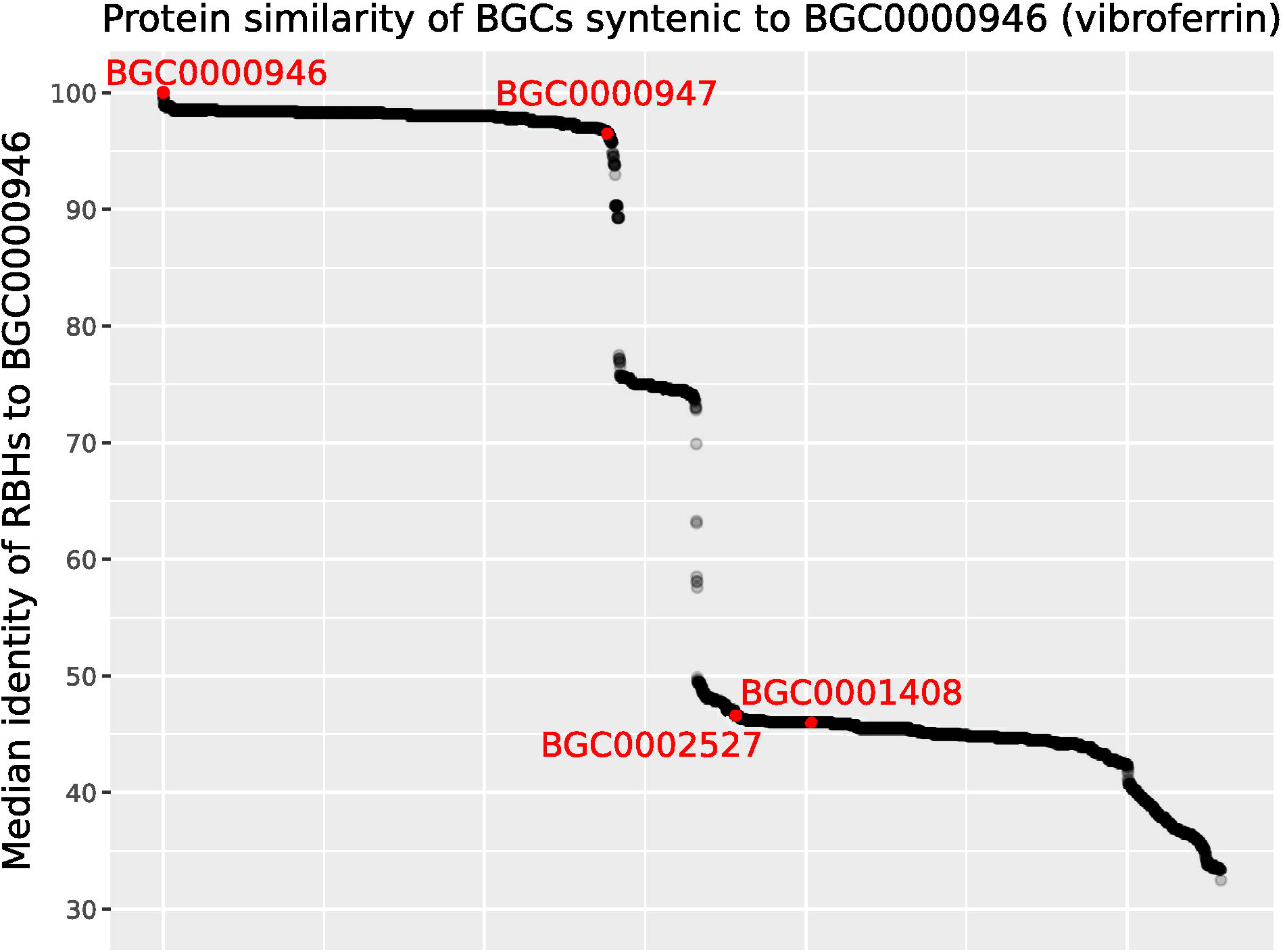
A SocialGene database containing more than 340,000 RefSeq genome assemblies and MIBiG BGCs was searched for gene regions complete and syntenic to MIBiG BGC000946 (encoding for vibrioferrin). The 6,581 points in the graph represent the resulting target BGCs and the y-axis represents the median protein identity of a BGC’s reciprocal best hits (RBHs) to BGC000946 proteins. Target BGCs were sorted in the x-axis by median RBHs to BGC000946 proteins. All MIBiG BGCs were labeled and highlighted in red and are known vibrioferrin producing BGCs, except BGC0001408 for which an associated chemical structure has not been reported.

Though tempting to believe the gradients would represent functional evolution and diversification of end-product SMs, one of the lowest median RBHs (46.6%) belonged to MIBiG BGC0002527, a vibrioferrin-producing BGC from *Azotobacter vinelandii* CA. The actual lowest median RBH of 32.5% was found in a *Facilibium subflavum* assembly. While there’s no evidence this *F. subflavum* strain produces the vibrioferrin siderophore, the flanking genes suggest the region is involved in metal acquisition and homeostasis (Supplementary Fig. 11).

While Fig. 4 shows 6,581 intact and syntenic BGCs, it is also possible these are situated within broader genomic contexts that catalyze the modification of vibrioferrin or an alike molecule. However, it is unclear what proportion of the BGCs this is, if any. Further studies are needed to determine the cause of the stepped gradients and whether they are due to speciation, horizontal transfer (see Supplementary Fig. 12), or other mechanisms. While outside the scope of the current study it is possible to create comprehensive in-database similarity links between BGCs for studying phylogenetic histories, especially those that are difficult to express as a phylogram.

### Extending capability by computing new nodes and relationships, in-database

SocialGene’s default schema is useful on its own but also designed to serve as a foundation from which new nodes and relationships can be created, a strength of Neo4j. For example, the SocialGene RefSeq database contains all MIBiG BGCs, antiSMASH predictions across 343,381 genomes, and MMseqs2 protein clustering to 90%, 70%, 50%, and 30% sequence identities. By traversing the existing nodes and relationships in the graph database a new type of relationship can be calculated that directly connects MIBiG BGCs to any genome assembly containing a similar BGC (Fig. 5). As shown in Fig. 5, the ability to filter subgraphs by additional metadata (e.g. taxon, host, etc.) enables researchers to create hypotheses about patterns of BGC distribution. Further, filtering or coloring availability in a culture collection provides a fast route to procuring strains for further experiments (Supplementary Fig. 14).

**Figure 5.**
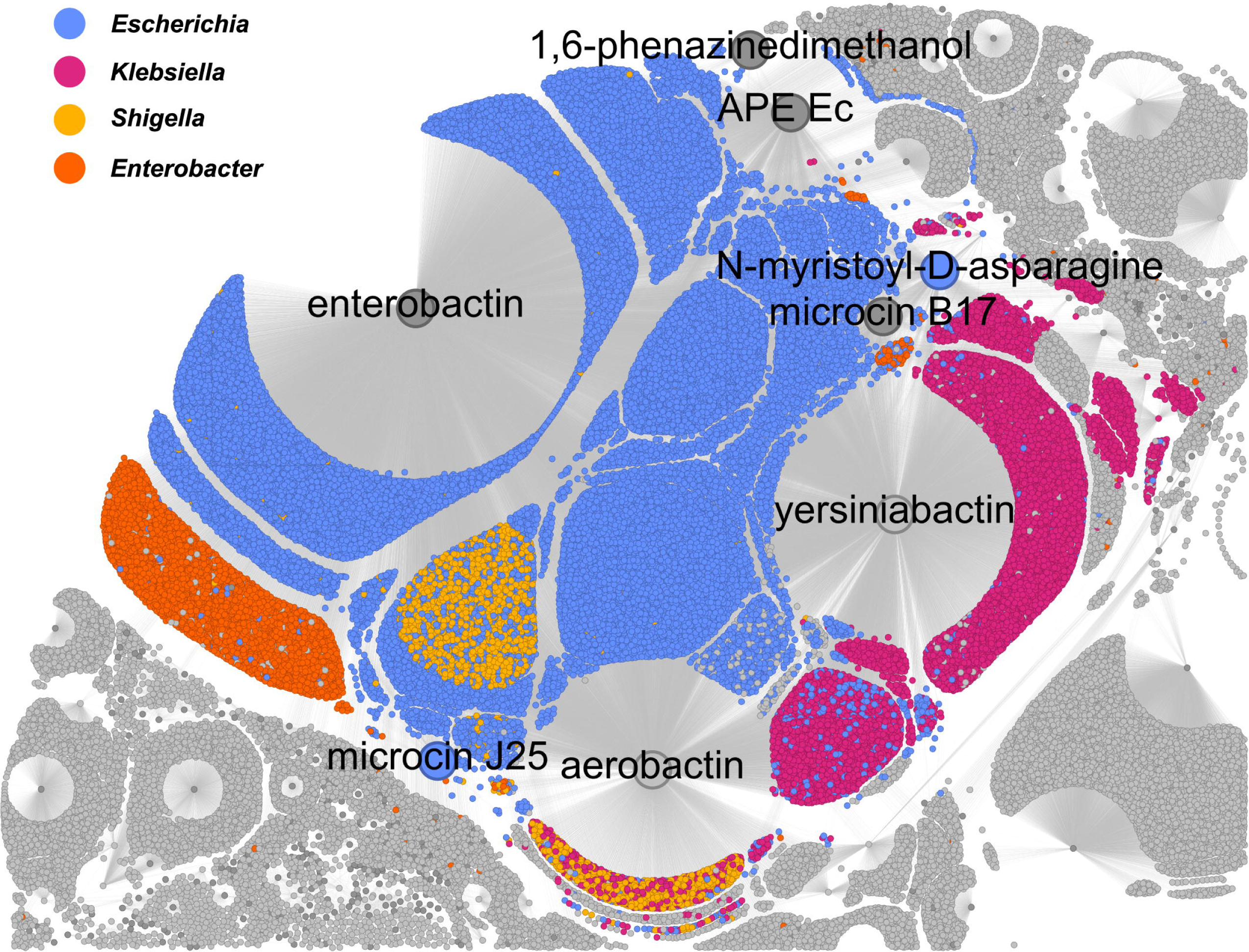
Using a single Neo4j Cypher statement, new links (edges between nodes) were created between MIBiG BGCs (nodes; some enlarged and labeled by SM product) and genomes assemblies (nodes). New links were created when an assembly contained an antiSMASH predicted BGC with proteins that were at least 70% similar to 70% of a MIBiG BGC’s proteins, as determined by traversing MMseqs2 protein cluster relationships. This figure is only a subset of the resulting subgraph (54% of total nodes) and highlights how SocialGene can be used to study complex distributions of BGCs, at scale. See Supplementary Figs. 13-15, for the full subgraph.

### Atlas of BGCs available in culture collections

One goal of creating the BGC search function in SocialGene was to enable repository-scale searches for BGCs across public and private culture collections. This aimed to uncover new sources of previously inaccessible BGCs, identify higher-yield strains, and provide a tool for hypothesis testing. However, we recognize that not everyone will have the necessary computational resources or expertise to install the SocialGene RefSeq database.

To address this, an online interactive atlas of MIBiG BGCs was developed, where similar BGCs can be found in various strain collections (e.g., NRRL, ATCC, DSMZ, etc.). This was achieved by searching each of the 2,502 MIBiG BGCs against the SocialGene RefSeq database, focusing specifically on the 27,406 genomes with metadata indicating availability in a culture collection. The resulting clustermap.js^59^ plots, restricted to 100 target BGCs per query MIBiG BGC (limited by visualization), include a total of 92,936 target BGCs spread across 2,112 MIBiG BGCs. For access to the atlas, refer to the Data and Code Availability section.

### Supporting narrow and broad meta-analyses

Version 3.0 of the MIBiG repository contains 2,502 BGCs and, like many natural product databases (e.g. npatlas^36,77^), entries are skewed towards well-studied taxa (e.g. Actinobacteria, especially *Streptomyces* spp.) and biosynthetic classes (e.g. PKS, NRPS, etc.). Despite this, early versions of MIBiG have been invaluable for building software and evaluating how computational methods and models behave with validated BGCs. SocialGene’s Nextflow workflow contains an optional flag (“--mibig”) which signals for the incorporation of all MIBIG BGCs into a SocialGene database, with or without additional input genomes.

For development and proof of concept work a SocialGene database was created containing all MIBiG BGCs. This resulted in a modest-sized graph database with 2.7 million nodes and 4.9 million edges, including more than 40,000 non-redundant protein nodes and more than 500,000 pHMM annotation relationships. Additionally, as many MIBiG BGCs contain NCBI taxonomy identifiers, SocialGene’s Nextflow “--ncbi_taxonomy” flag was used, which downloads and parses the entire NCBI taxonomy database, and links input BGCs/genomes to the source organism in the taxonomy graph. Supplementary Fig. 16 visualizes this placement of all MIBiG BGCs onto the taxonomic graph within a SocialGene database and highlights the taxonomic bias. We also exported a subgraph of all non-redundant proteins, less-redundant pHMMs, and the annotation links connecting the two, for import and layout in Gephi (Supplementary Fig. 17).

As expected, proteins were primarily clustered by function, but the graph also excelled in displaying the complicated evolutionary relationships between both large multidomain and smaller accessory proteins. Similar analysis allows for putative functional transfer to hypothetical proteins.

### Targeted antibiotic drug discovery

Intentional query engineering leveraging *in silico*, *in vitro*, and *in vivo* domain knowledge enables targeted large-scale searches of SocialGene databases across biochemistry, chemistry, and modes of action. These targeted searches and analyses can be designed and used to inform wet-lab experiments pre-, ad-, and post hoc. For example, a customized search for peptidic and halogenated antibiotics can guide the choice of isolation and bioassay techniques.

To engineer such a search we can exploit the fact that bacteria often encode resistance mechanisms within a BGC to counteract the toxicity of the produced specialized metabolite(s). Searching for self resistance proteins is a strategy incorporated into other genome mining software such as ARTS.^78^ As protein functional information is present in SocialGene, in the form of pHMM annotations, similar strategies can be employed in-database without prior BGC detection. Using a similar strategy we can also detect putative halogenase and NRPS enzymes and their co-occurrence.

For example, Supplementary Fig. 18 displays a query for any nucleotide sequence containing a protein annotated by a tryptophan halogenase pHMM, within 10 kb of a nonribosomal peptide synthetase (NRPS) protein (detected with antiSMASH’s pHMM detection rule, performed in-database), and within 50 kb of a protein annotated by an AMRfinder^48,79,80^ pHMM (antimicrobial resistance gene detection). When limited to MIBiG sequences, the query only returned halogenated NRP antibiotics such as vancomycin (Supplementary Fig. 19). Because chlorinated and brominated natural products provide characteristic isotopologue distributions, peptidic SM often fragment well in ESI LC-MS/MS, and antibiotic activity is suggested, subsequent lab work can be intentional and directed.

### Targeted drug discovery

SocialGene facilitates targeted and untargeted drug discovery beyond microbial antibiotics. For instance, Pfam PF00227 is a multi-kingdom pHMM model of the "proteasome subunit". Proteasomes are ancient multi-subunit proteases^81,82^ involved in controlled protein degradation and recycling, and small molecule inhibitors targeting proteasomes have provided promising candidates for cancer therapeutics.^83,84^ As proof of concept, the SocialGene RefSeq database was searched for MIBiG BGCs containing a protein annotated by the Pfam pHMM model PF00227, of which there were eight. All eight of the BGCS produce proteasome inhibitors: fellutamide B^85^, cinnabaramide A^86^, landepoxcin^87^, salinosporamide A^88–90^ (two BGCs), clarexpoxcin^87^, eponemycin^91–93^, and TMC-86A^94^. Next, an identical search was performed but restricted instead to the over two million antiSMASH-predicted BGC regions in the RefSeq genomes. While the MIBiG BGCs consisted of PKS, NRPS, and PKS/NRPS hybrids, this larger search revealed 1,595 diverse BGCs from 25 phyla across Eukaryota, Archaea, and Bacteria. BGC type counts are displayed in Supplementary Table 1. Enabling fast searches (the above search takes milliseconds) allows users to quickly iterate over potential targets and hypotheses.

### Restricting pHMM annotations to specific motifs

While pHMM annotations provide a rapid means for discovering proteins with specific functions they may prove too general for some tasks. However, highly targeted searches can be performed using in-database regex filtering of the non-redundant protein amino acid sequences, either alone, or in addition to pHMM annotation queries.

For example, proteins in the CAP protein superfamily^95^ family are implicated in various biological roles, including virulence and facilitating host-symbiont/pathogen relationships^96^. Based on this, one could hypothesize that BGCs containing a CAP might also be involved in virulence and/or cross-kingdom interactions. However, across the 343,381 genomes in the SocialGene RefSeq database, 135,595 genomes contain a protein(s) annotated by Pfam pHMM PF00188 (CAP superfamily), and in 5,593 of those a CAP is in an antiSMASH 7-predicted BGC. As this is still an unmanageable number, we could further restrict our hypothesis. For example, Hirano et al. recently suggested that insect cysteine-rich secretory proteins (CAPs) may induce the formation of plant galls^97^ and, specifically, CAPs with the core sequence (F/Y-T-Q-I/V-V-W), which can be expressed with the regex pattern ".*[FY]TQ[IV]VW.*". Filtering on PF00188 and this pattern reduced the number of genomes to 2,337; and limiting the results to antiSMASH 7-annotated regions returned a reasonable 43 genomes (Supplementary Fig. 20). Each of these searches completed in less than one second, which allows fast iteration over hypotheses.

### Extending SocialGene

It is easy to extend the current graph schema both at database creation and within an active database. Neo4j has a number of import and cross-database integration tools, including being able to read data directly from web-hosted SPARQL endpoints. This provides the opportunity for future integration of additional data from sources such as ChEMBL^98^, UniProt^45^, etc. Additionally, the Python library and Nextflow workflows were written modularly to allow add-ons of new node and relationship types.

### Connecting chemistry to biology

Socialgene’s Python library has an add-on that parses and integrates nodes and relationships representing the Natural Products Atlas (NP Atlas) into an active SocialGene database (Supplementary Fig. 21). Many NP Atlas chemical structures are linked to a species-level NCBI taxonomic identifier of the organism from which the compound was first reported. However, species level taxonomic IDs are often not specific enough to correlate a chemical compound with the genome assembly of a specific producer. To alleviate this, the add-on creates an additional link using simple text-similarity measures on taxa names between NP Atlas and NCBI taxonomy. In addition to including NP Atlas metadata, SocialGene creates non-redundant chemical nodes within the graph database which are linked by Tanimoto similarity (Supplementary Figs. 22, <u>23</u>).

For users with paired genome and mass spectrometry data, GNPS networking^37^ results can be downloaded and integrated directly. For example, Supplementary Fig. 24 shows the ingestion and incorporation of the GNPS molecular networking and library search results from 122 LC-MS/MS runs across 120 bacterial isolates (data previously published^99^). Additionally, Supplementary Figs. 25, <u>26</u> show the resulting interconnections between genomes; mass spectra; GNPS clusters, networks and library hits; and NP Atlas entries. Future research will aim to build models correlating MS features, clusters, and chemical moieties to BGCs, sub-BGCs, proteins, and protein domains.

### Limitations and improvements

Machine learning/artificial intelligence for protein function inference has significantly advanced during the years-long creation of SocialGene, including AlphaFold protein structure prediction^71^ and the subsequent dramatic increase in available predicted structures.^100^ While future additions to SocialGene should include methods such as Foldseek^101^ (combination of protein sequence and structure alignment) it would rely on input sequences being present in the DeepMind/EMBL-EBI’s AlphaFold Protein Structure Database or running AlphaFold. The latter would require an increase in SocialGene’s complexity and compute requirements and including full models would require hundreds of additional gigabytes of storage, even with compression.^102^ Additionally, the AlphaFold Protein Structure Database is currently limited to proteins less than 2,700 AA for proteomes/Swiss-Prot and 1,280 AA for the rest of UniProt^46^. Another promising avenue, with a preliminary analysis by Schütze et al.^103^ is to utilize the vector similarities of protein model embeddings directly. This is possible in the latest versions of Neo4j, which incorporate vector indexing, cosine similarity, and K-Nearest Neighbors searches. However, as discussed by Schütze et al,^103^ and our own experimentation, the speed of generating embeddings can be slow for large scale experiments, especially on commodity hardware. UniProt currently distributes embeddings generated from prottrans_t5_xl_u50, though this only covers UniProtKB/Swiss-Prot^46^; however, it may be feasible to include publicly released ESM-2 embeddings^104^ in the future.

In the SocialGene RefSeq database 81% of the more than 304 million proteins are annotated by at least one pHMM model, and 79.4% by Pfam models alone; this is similar to a report in 2019 that 77.2% of UniprotKB proteins had at least one Pfam annotation.^105^ Along with greater pHMM coverage, SocialGene could be improved by community analysis and clustering of proteins by domain architecture in-database, or calculated externally using tools such as DAMA.^106^

Apart from MIBiG, the datasets in this manuscript were created from all, or subsets, of RefSeq. RefSeq was chosen for proof-of-concept as it had the highest number of publicly available genomes with mostly-consistent genome annotation.^41^ However, there are a number of limitations with applying at-scale analyses on current public databases, including RefSeq. We especially caution anyone using our premade RefSeq databases, or other non-curated genomes, for ecology and evolution studies. For such studies SocialGene has the ability to use nucleotide FASTA files as input to the Nextflow workflow, which are then analyzed with Prokka^107^ for consistent gene annotation; or users can provide their own gene-called genomes.

### Cost to create a database

Nextflow Tower was used to run the SocialGene Nextflow workflow on 314 *Micromonospora* genomes. This included the compute-intensive steps of: annotating proteins with antiSMASH^33^, AMRFinder^48^, Pfam^50^, Resfams^52^, and TIGRFAM^53^ pHMMs; clustering proteins to three levels with MMseqs2; and running antiSMASH v7^33^ on all 314 genomes. Running the full workflow on AWS Batch cost less than $5.00 USD, unoptimized. However, with the ability to run multiple analyses on large amounts of data, we suggest users using fee-based computing resources conduct their own cost estimate experiments before scaling.

## Data and Code Availability

SocialGene’s source code is hosted at <u>github.com/SocialGene</u>, with documentation at <u>socialgene.github.io</u>. The Python library is available at <u>github.com/socialgene/sgpy</u> and doi.org/1<u>0.5281/zenodo.12207092</u>; the Nextflow workflow is available at <u>github.com/socialgene/sgnf</u> and doi.org/1<u>0.5281/zenodo.12207039</u>. Code and notebooks used to generate manuscript figures and analyses are available at <u>github.com/socialgene/manuscript_1</u> and doi.org/1<u>0.5281/zenodo.13333842</u>. All SocialGene databases computed for this manuscript are archived as Neo4j dumps at: doi.org/<u>10.5061/dryad.ns1rn8q2k</u>. The BGC atlas is available interactively at <u>bgcatlas.pages.dev</u> and archived at doi.org/<u>10.5281/zenodo.12775149</u>.

## Acknowledgements

This research was supported by an NIGMS grant R35GM133776. C.M.C was additionally supported by an NLM training grant to the Computation and Informatics in Biology and Medicine Training Program (NLM 5T15LM007359). Portions of this research was performed using the compute resources and assistance of the UW-Madison Center For High Throughput Computing (CHTC) in the Department of Computer Sciences. The CHTC is supported by UW-Madison, the Advanced Computing Initiative, the Wisconsin Alumni Research Foundation, the Wisconsin Institutes for Discovery, and the National Science Foundation, and is an active member of the OSG Consortium, which is supported by the National Science Foundation and the U.S. Department of Energy’s Office of Science. Portions of this research was done using services provided by the OSG Consortium,^108,109^ which is supported by the National Science Foundation awards #2030508 and #1836650.

## Author Contributions

C.M.C. Conceptualization, Data curation, Formal Analysis, Investigation, Methodology, Visualization, Writing – original draft, Writing – review & editing, Funding acquisition. JCK: Conceptualization, Editing, Supervision, Funding acquisition

