## Supplementary Information for "Creating and leveraging bespoke large-scale knowledge graphs for comparative genomics and multi-omics drug discovery with SocialGene"

Chase M. Clark[1], Jason C. Kwan[1*]
[1]Division of Pharmaceutical Sciences, School of Pharmacy, University of Wisconsin-Madison, 777 Highland Avenue, Madison, WI 53705, USA

### Supplementary Figure 1: Simplified outline of data processing

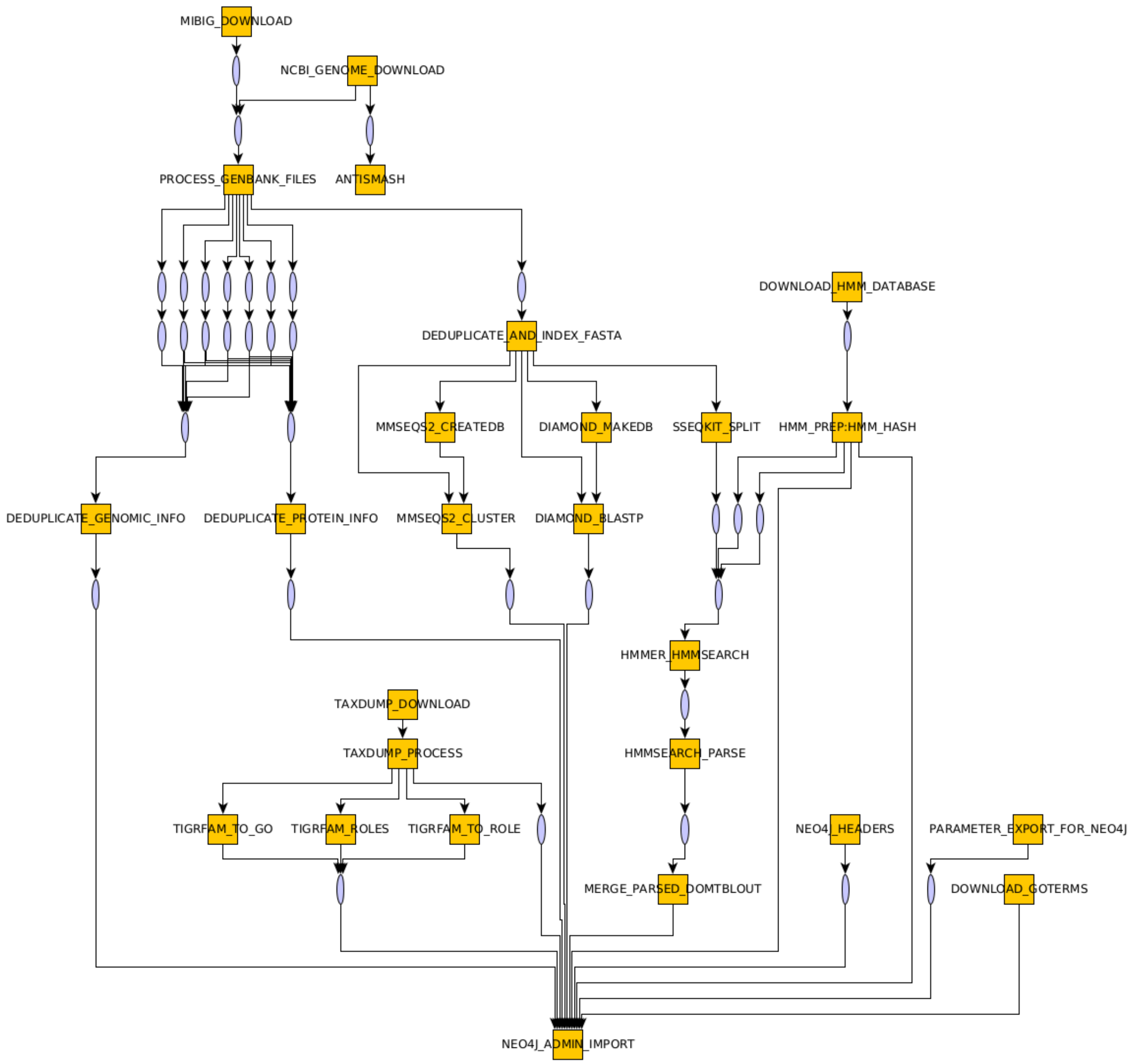


This outline shows a simplified outline of the automated steps taken by SocialGene's Python and Nextflow libraries to take input genomes and transform them into a Neo4j database. Square boxes represent an action or actions and ovals represent a file/unit of data.

### Supplementary Table 1: SocialGene RefSeq database

| node label | count |
|---|---|
| amrfinder | 630 |
| antismash | 940 |
| assembly | 345883 |
| culture_collection | 152 |
| goterm | 51283 |
| hmm_source | 25813 |
| hmm | 25566 |
| nucleotide | 46179608 |
| parameters | 1 |
| pfam | 19632 |
| protein | 304330794 |
| resfams | 123 |
| taxid | 2543881 |
| tigrfam_mainrole | 19 |
| tigrfam_role | 116 |
| tigrfam_subrole | 101 |
| tigrfam | 4488 |

| relationship type | count |
|---|---|
| ALTERNATE | 3646 |
| ANNOTATES | 847850795 |
| ASSEMBLES_TO | 46179608 |
| ENCODES | 1337412265 |
| FOUND_IN | 27874 |
| GO_ANN | 6387 |
| IS_A | 28045 |
| IS_TAXON | 343763 |
| MAINROLE_ANN | 116 |
| MMSEQS_30 | 27681483 |
| MMSEQS_50 | 54434786 |
| MMSEQS_70 | 111454401 |
| MMSEQS_90 | 304330794 |
| NEGATIVELY_REGULATES | 2684 |
| PART_OF | 6323 |
| POSITIVELY_REGULATES | 2682 |
| PROTEIN_TO_GO | 35527 |
| REGULATES | 3100 |
| ROLE_ANN | 3169 |
| SOURCE_DB | 25813 |
| SUBROLE_ANN | 116 |
| TAXON_PARENT | 2543881 |

The number of nodes and relationships in the base SocialGene RefSeq database. This includes the 343,381 RefSeq genomes along with 2,502 MIBiG BGCs (included under "assembly" and "nucleotide" nodes). Note that these only represent the database as built by the Nextflow pipeline. Additional nodes and relationships mentioned in the manuscript were added through use of add-ons to the SocialGene Python library as they are not suited for import via the Neo4j admin-import command, as they require a running database to make connections to existing nodes.

### Supplementary Figure 2: Outdegree distribution of pHMM nodes to non-redundant protein nodes

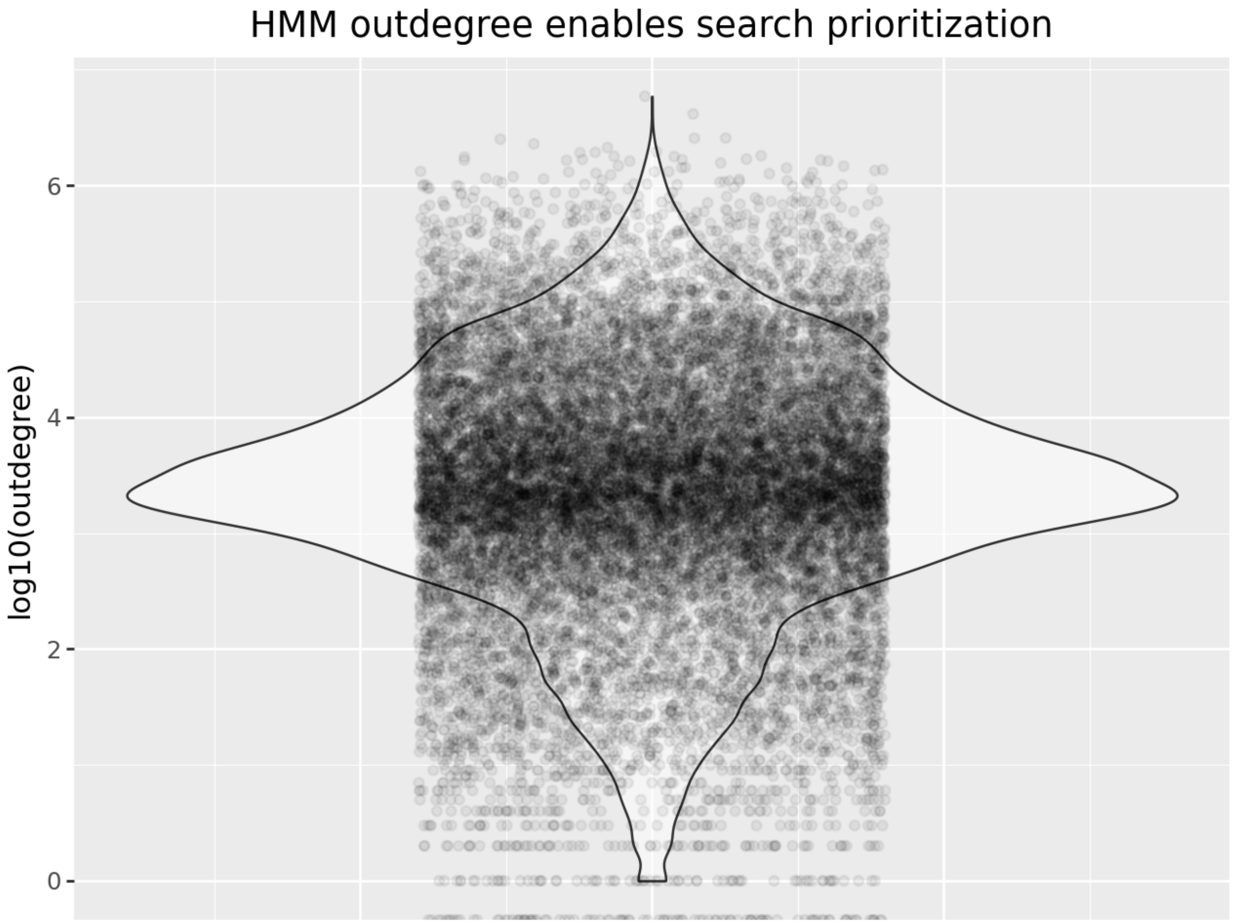


When searching a SocialGene database for similar BGCs one of the prioritization methods within SocialGene's search function is to first rank the pHMM models that annotate the query BGC's proteins based on the number of "ANNOTATES" relationships each of the pHMM model nodes has to non-redundant protein nodes. Prioritizing pHMM models with a lower outdegree reduces the search space and has the side effect that the query domains and proteins that are prioritized are often less common. The figure shows the density of the log10 outdegree of the pHMM models and the 847,850,795 "ANNOTATES" relationships in the SocialGene RefSeq database.

### Supplementary Text 1

With a query BGC containing 20 proteins, the algorithm will find all pHMM models annotating the 20 proteins. The proteins are ranked by the ascending sum of their pHMM outdegrees and then a subset of the proteins used for the search. The number of proteins and the number of pHMM models to consider per protein are user-selected variables that are set as absolute or percent values. This allows for a faster search that prioritizes proteins with less frequently seen pHMM annotations and ignores ubiquitous domains and proteins, but can lead to missed BGCs depending on settings and which proteins are present in the query and target BGCs. For example, MIBiG entry BGC0001848 contains 50 proteins and is found in a *Micromonospora* sp. Searching >340,000 genomes for similar BGCs using the top 5 prioritized proteins, as described above, finishes in 2.5 seconds and recovers BGC0001848 and its originating RefSeq genome, among other closely-related BGCs. However, it fails to capture BGC0001850, which is encoded in a *Streptomyces* sp. and produces a similar compound, but only shares 20 of BGC0001848's 50 proteins in a different genetic arrangement; see Braesel et al[1] for detailed analysis of these two clusters. The search function provides the ability to bypass the filters outright, or selectively, for defined query proteins. Future efforts will go towards balancing this search algorithm's efficiency and accuracy.

#### Supplementary Figure 3: Within-cluster comparison of DIAMOND BLASTp and MMseqs2

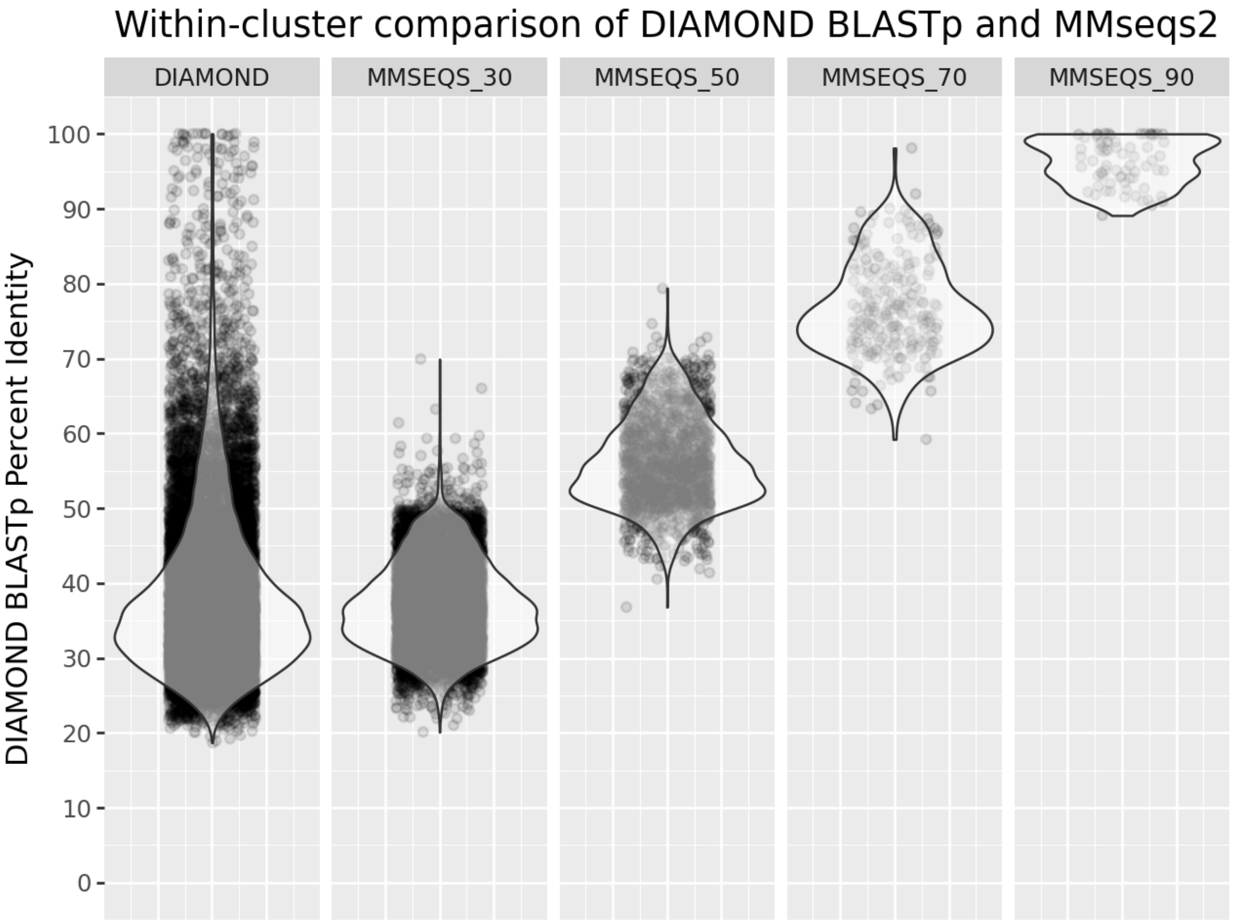


While the MMseqs2 clustering in SocialGene isn't perfect (possibly due to cascaded clustering being run in SocialGene without cluster reassignment), the MMseqs2 relationships within SocialGene all fall within tolerance of their value (i.e. MMSEQS_50 relationships usually fall within 50%-70% identity. Each point in the figure represents a DIAMOND BLASTp percenet identity score between two within-cluster proteins (e.g. two proteins within the same MMSEQS_90 cluster).

Data shown is from a SocialGene database created with genomes GCF_000009045.1, GCF_000005845.2, GCF_008931305.1 (*Bacillus subtilis* subsp. *subtilis* str. 168, *Escherichia coli* str. K-12 substr. MG1655, and *Streptomyces coelicolor* A3(2); downloaded on June 20, 2024); MMseqs2 clustering in steps of 90, 70, 50, 30 with settings '-c 0.7 --cov-mode 0 --split 1'; all-vs-all DIAMOND BLASTp was run with settings '-k0 --max-hsps 1 --query-cover 70 --subject-cover 70 --block-size 6'.

#### Supplementary Figure 4: Inter-cluster comparisons of DIAMOND BLASTp and MMseqs2

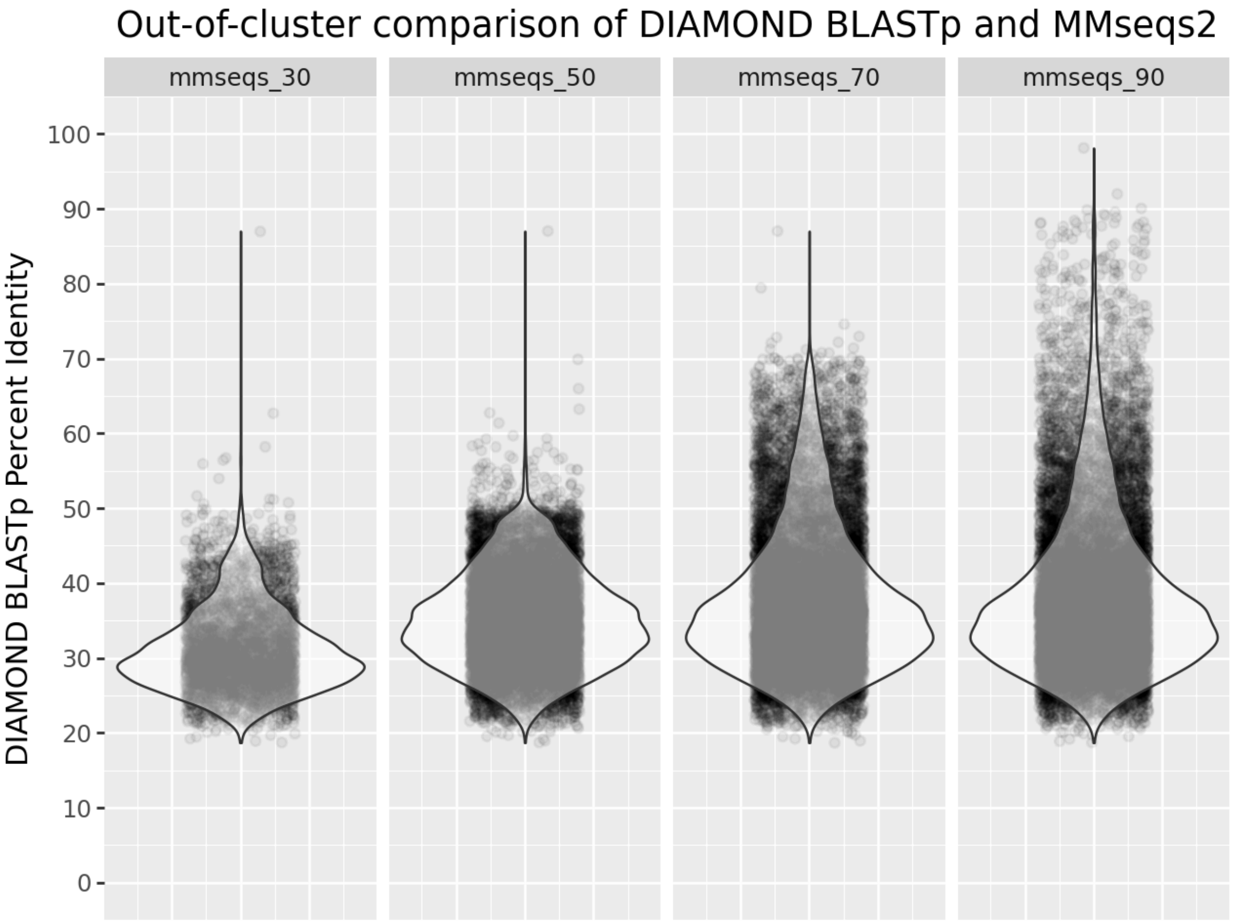


The inter-cluster similarities can sometimes be higher than the MMseqs2 clustering level (e.g. two separate 30% clusters may contain member(s) that share >30%) and should be considered when designing queries/experiments. Each point in the figure represents a DIAMOND BLASTp comparison between two inter-cluster proteins. For example, a point in the 'mmesqs_90' column represents the DIAMOND BLASTp identity of two proteins belonging to different 'mmesqs_90' clusters.

Data shown is from a SocialGene database created with genomes GCF_000009045.1, GCF_000005845.2, GCF_008931305.1 (*Bacillus subtilis* subsp. *subtilis* str. 168, *Escherichia coli* str. K-12 substr. MG1655, and *Streptomyces coelicolor* A3(2); downloaded on June 20, 2024); MMseqs2 clustering in steps of 90,70,50, 30 and settings '-c 0.7 --cov-mode 0 --split 1'; all-vs-all DIAMOND BLASTp was run with settings '-k0 --max-hsps 1 --query-cover 70 --subject-cover 70 --block-size 6'.

#### Supplementary Figure 5: Lack of correlation between SocialGene's domain similarity and DIAMOND BLASTp identity

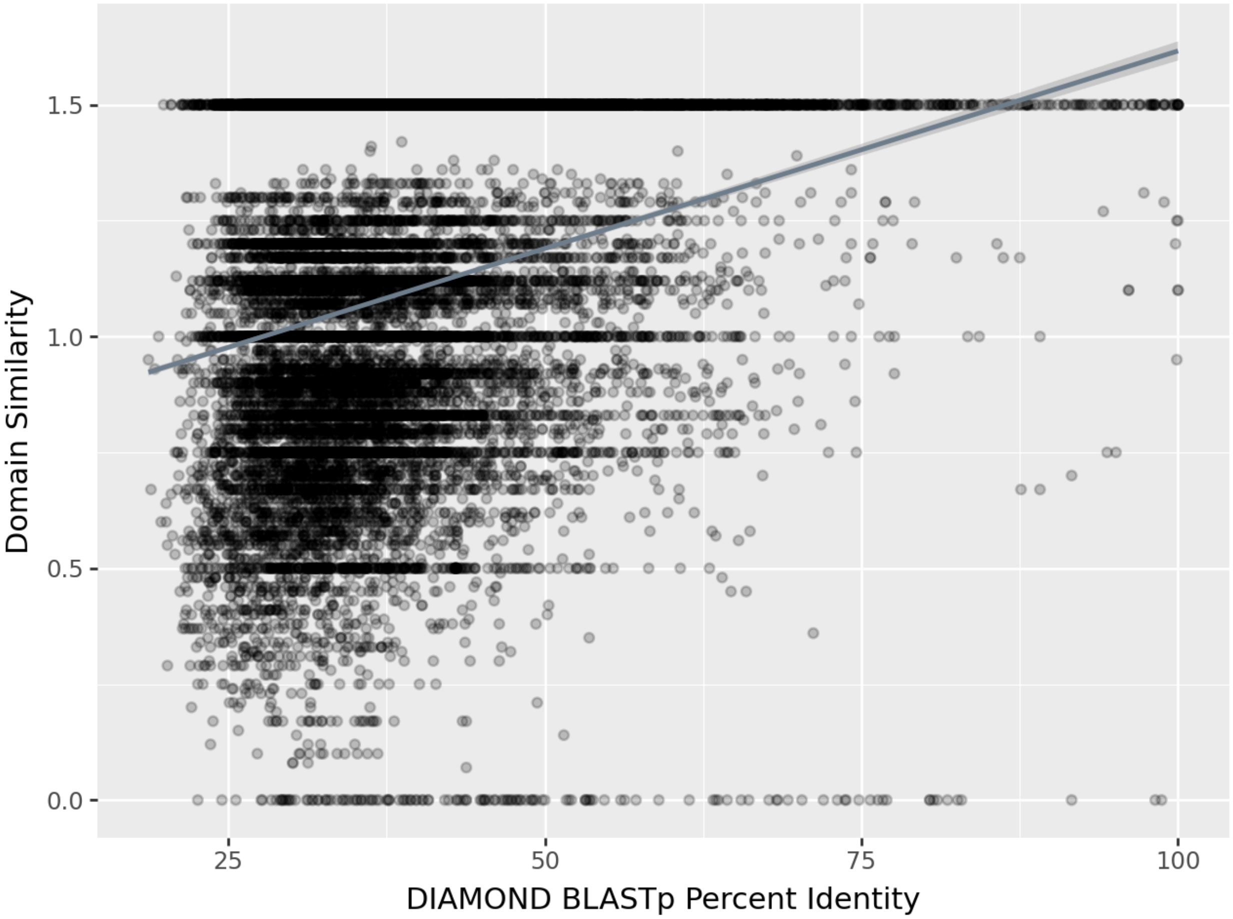


Domain similarity, measured using a combined, weighted Levenshtein and Jaccard similarity score, shows a lack of correlation with sequence alignment scores. This finding is discussed further in the manuscript.
Linear regression: slope 8.6; intercept 29.0; rvalue 0.27; pvalue 0.0; stderr 0.17
Data shown is from a SocialGene database created with genomes GCF_000009045.1, GCF_000005845.2, GCF_008931305.1 (*Bacillus subtilis* subsp. *subtilis* str. 168, *Escherichia coli* str. K-12 substr. MG1655, and *Streptomyces coelicolor* A3(2); downloaded on June 20, 2024); all-vs-all DIAMOND BLASTp was run with settings '-k0 --max-hsps 1 --query-cover 70 --subject-cover 70 --block-size 6'; pHMM databases included "antismash, amrfinder, pfam, resfams, tigrfam".

#### Supplementary Figure 6: 3D alignment of AlphaFold predicted protein structures

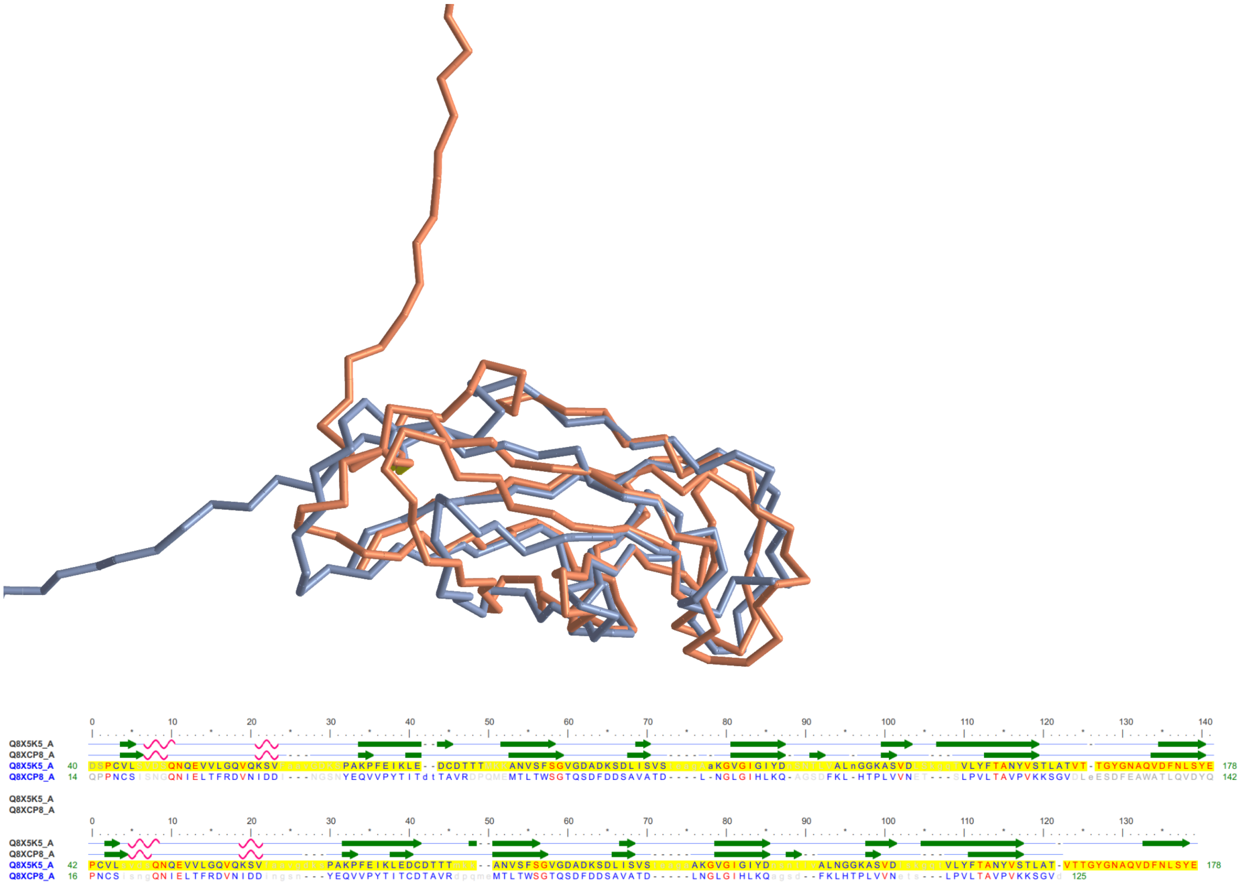


Alignment of the AlphaFold[2] predicted 3D protein structures (AlphaFold DB version 2022-11-01) using TM-align[3] through NCBI-hosted iCn3D[4,5] showed that, while the proteins have highly dissimilar AA sequences, they have similar predicted 3D structures (RMSD: 2.487 Å, TM-score: 0.7134; approximately 15-20% of the proteins' residues remain unfolded).
Structures for proteins Q8X5K5 (orange, with unfolded region pointing up) and Q8XCP8 (blue, with unfolded region pointing left). The two proteins share such low sequence identity that they are not aligned by BLASTp's default settings, but are both annotated by a single PFAM pHMM, PF00419.23. The aligned, predicted 3D protein structures can be viewed at:
https://structure.ncbi.nlm.nih.gov/icn3d/share.html?Ei8f9WLdzgqaRQXv7

### Supplementary Text 2:

A nearly-perfect BLASTp alignment doesn't necessitate similar pHMM annotation. For example, UniProtKB A0A0H3JI96 and A0A0H3JGM8 are phage tail proteins encoded in the *Escherichia coli* O157:H7 genome. While BLASTp alignment revealed matches in 233 of 238 positions (97.9% identity), only a third of their pHMM annotations overlapped. Inspecting the PFAM pHMM annotations for these proteins through UniProt's web service (on 05-01-2023) revealed A0A0H3JGM8 was annotated by model PF13927 and A0A0H3JI96 by a different model, PF07679. However, aligning PFAM models with pHMMscan using European Bioinformatics Institute's (EBI's) pHMMerWeb version 2.41.2 resulted in A0A0H3JGM8 being annotated by models PF13927, PF13895, and PF07679 (though only PF13927 was reported, as it was chosen to represent the PFAM clan) and A0A0H3JI96 being annotated by model PF07679. The discrepancy between UniProt and EBI's annotation of A0A0H3JGM8 was likely due to different post-processing in choosing PFAM clan members. Currently SocialGene doesn't prune PFAM pHMM annotations based on PFAM clan membership and so it reported A0A0H3JGM8 as annotated by PF13927, PF13895, and PF07679; and A0A0H3JI96 as annotated by PF07679, resulting in them sharing a single annotation, of three, for a similarity score of 0.33. Future versions of SocialGene may work to address a reproducible method for pruning PFAM annotations by clan. Nevertheless, it remains that pHMM annotation can be disrupted even at high BLASTp identities and, at the time of writing, comparing PFAM annotations between any of EBI's pHMMerWeb, UniProt, and SocialGene could lead to missing proteins with similar annotations. In this case, comparing UniProt annotations to EBI's pHMMerWeb would have resulted in missing the similar proteins, whereas SocialGene didn't, due to consistent pHMM annotation.

### Supplementary Figure 7: Recovery of a known lagriamide B BGC from *Paraburkholderia acidicola*, using the Lagriamide A producing BGC (BGC0001646) as the search query

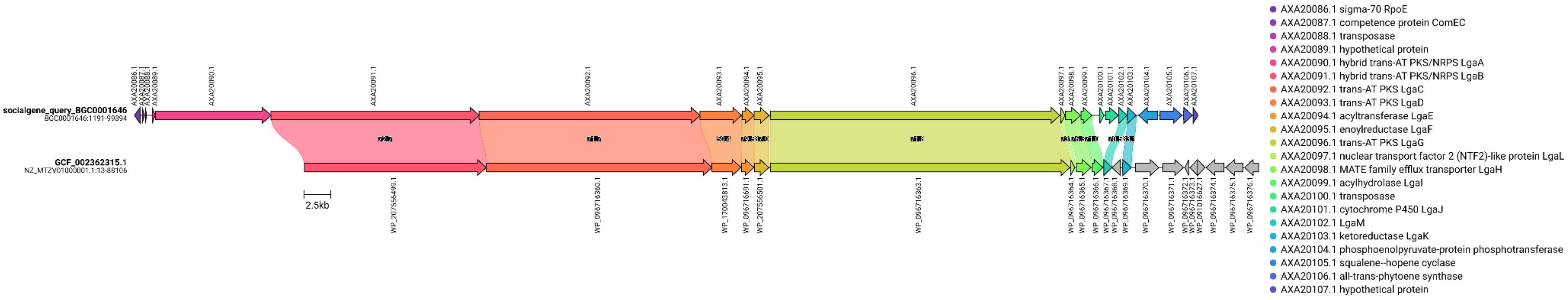


The SocialGene BGC search was run against the RefSeq SocialGene database using the following parameters: use_neo4j_precalc: True; assemblies_must_have_x_matches: 0.6; nucleotide_sequences_must_have_x_matches: 0.6; gene_clusters_must_have_x_matches: 0.6; break_bgc_on_gap_of: 20000; target_bgc_padding: 10000; max_domains_per_protein: 3; max_outdegree: 300000; max_query_proteins: 10; scatter: True; locus_tag_bypass_list: None; protein_id_bypass_list: None; only_culture_collection: False; frac: 0.75; run_async: True; analyze_with: "blastp"

Additional partial match, lower-scoring, target BGCs were removed for figure clarity. Both a high resolution and interactive version of the plot are available in the archived manuscript's repository (see manuscript's Data and Code Availability section).

### Supplementary Figure 8: Fragmented rapamycin BGCs found by SocialGene BGC search

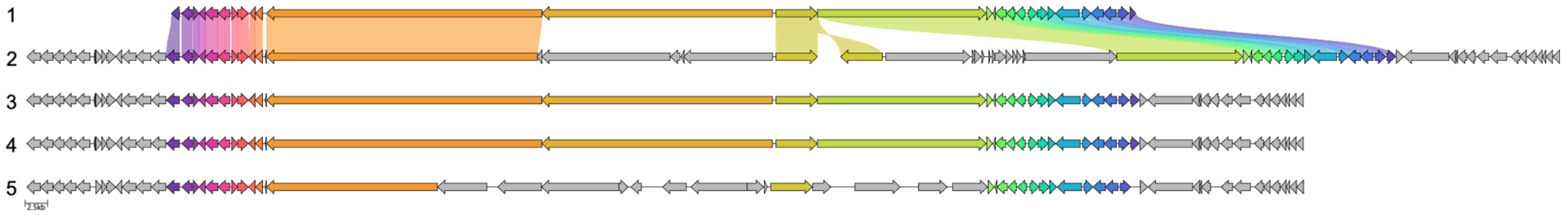


| | Assembly | Org | Sequencing | Assembly | Gene Calling |
|---|---|---|---|---|---|
| **1** | BGC0001040 | University of Cambridge | ABI 373A | Staden Package | BLAST/FASTA |
| **2** | GCF_014205315.1 | DOE Joint Genome Institute | PacBio | Flye v. 2.6 | PGAP 6.6 |
| **3** | GCF_003675955.1 | Novartis AG | PacBio RSII | HGAP v. DEC-2014 | PGAP 6.4 |
| **4** | GCF_024298965.1 | Korea University | PacBio RSII; Illumina NovaSeq | HGAP v. 2.3 | PGAP 6.4 |
| **5** | GCF_000418455.1 | University of Kaiserslautern | 454 | GS De Novo Assembler (v 2.8) program (Roche) v. 2.8 | PGAP 6.4 |

All five of the above sequences derive from separate sequencing efforts of the same type strain, *Streptomyces rapamycinicus* DSM 41530/NRRL 549. Despite being the same organism, the rapamycin BGC in assembly 2 is split across two contigs and has fragmented polyketide synthase genes. Additionally, assembly 5 contains fragmented polyketide synthase genes. Despite this, SocialGene's search function was able to recover these BGCs when BGC0001040 was used as input.

The search was run against the RefSeq SocialGene database using the following parameters: use_neo4j_precalc=True; assemblies_must_have_x_matches=0.4; nucleotide_sequences_must_have_x_matches=0.4; gene_clusters_must_have_x_matches=0.4; break_bgc_on_gap_of=20000; target_bgc_padding=10000; max_domains_per_protein=3; max_outdegree=1000000; max_query_proteins=5; scatter=False; locus_tag_bypass_list=None; protein_id_bypass_list=None; only_culture_collection=False; frac=0.75; run_async=True; analyze_with="blastp"

There were additional assemblies with results but which were removed for figure clarity.

### Supplementary Figure 9: SocialGene BGC search recovering multiple BGC integrations

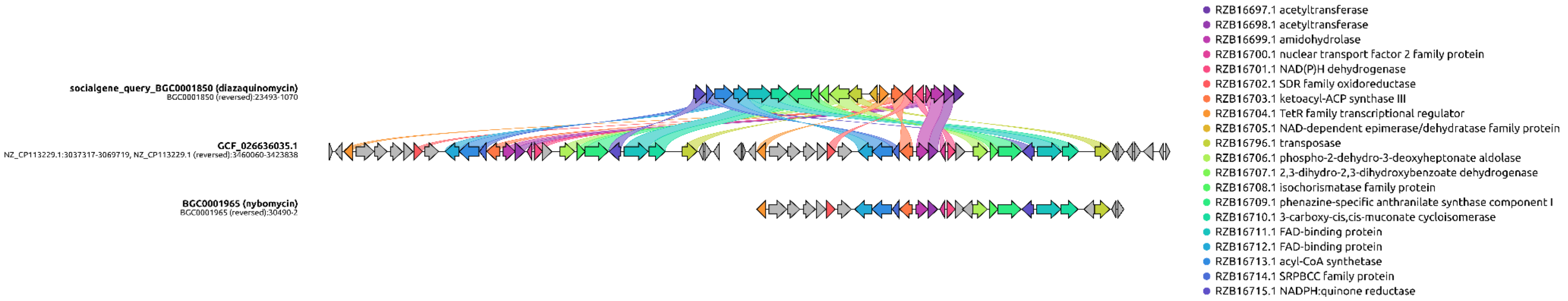


While using BGC0001850 (first BGC in figure, encodes for diazaquinomycin) to spot-check the results of querying all MIBiG BGCs against the RefSeq SocialGene database it was noticed that two assemblies had multiple copies of a similar BGC to BGC0001850. These BGCs were most similar to another MIBiG entry, BGC0001965 (bottom BGC in figure), which encodes for nybomycin, a chemical analog of diazaquinomycin. Further inquiry revealed the genome assembly GCF_026636035.1 to be a *Streptomyces albidoflavus* strain engineered for the overproduction of nybomycin[6] (middle BGC in figure). The second assembly recovered (not shown) was GCF_026210455.1, the BAC 4N24 plasmid donor from the same study [6].

The search was run against the RefSeq SocialGene database using the following parameters:
use_neo4j_precalc: True; assemblies_must_have_x_matches: 0.7; nucleotide_sequences_must_have_x_matches: 0.7; gene_clusters_must_have_x_matches: 0.7; break_bgc_on_gap_of: 10000; target_bgc_padding: 20000; max_domains_per_protein: 3; max_outdegree: 1000000; max_query_proteins: 10; scatter: False; locus_tag_bypass_list: None; protein_id_bypass_list: None; only_culture_collection: False; frac: 0.75; run_async: True; analyze_with: 'blastp'

There were additional assemblies with results but which were removed for figure clarity.

### Supplementary Figure 10: Putative pseudomonic acid BGCs

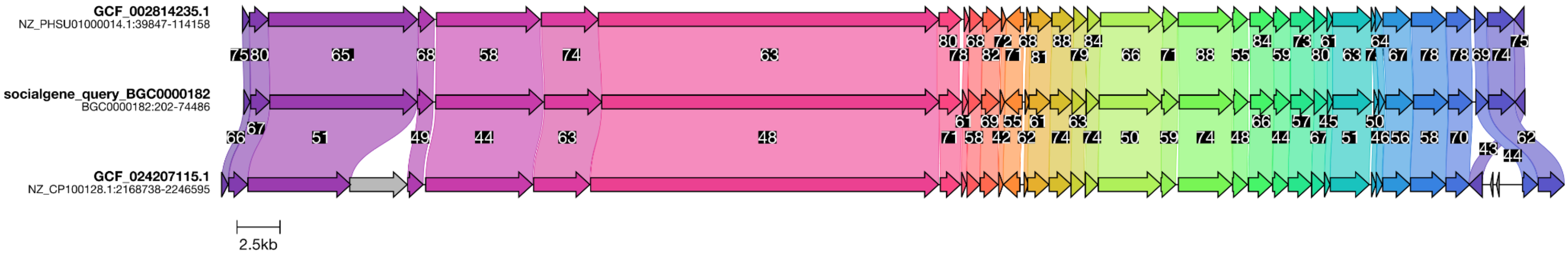


MIBiG BGC0000182 is a pseudomonic acid BGC from *Pseudomonas fluorescens*. When using SocialGene to search for related BGCs in the RefSeq SocialGene database two genomes had BGCs highly syntenic to BGC0000182 but with relatively low individual protein identities (black labels between homologous BGC genes). GCF_002814235 is *Pseudomonas* sp. QS1027 and GCF_024207115 is *Chromobacterium* sp. IIBBL 290-4 which falls in a separate taxonomic class. Both a high resolution and interactive version of the plot are available in the archived manuscript's repository (see manuscript's Data and Code Availability section).

The *Chromobacterium* BGC is available in antiSMASH DB[7]:
https://antismash-db.secondarymetabolites.org/output/GCF_024207115.1/index.html#r1c3

#### Supplementary Figure 11: Target BGC with the lowest median BLASTp similarity (34.1) to the vibrioferrin BGC, BGC0000946

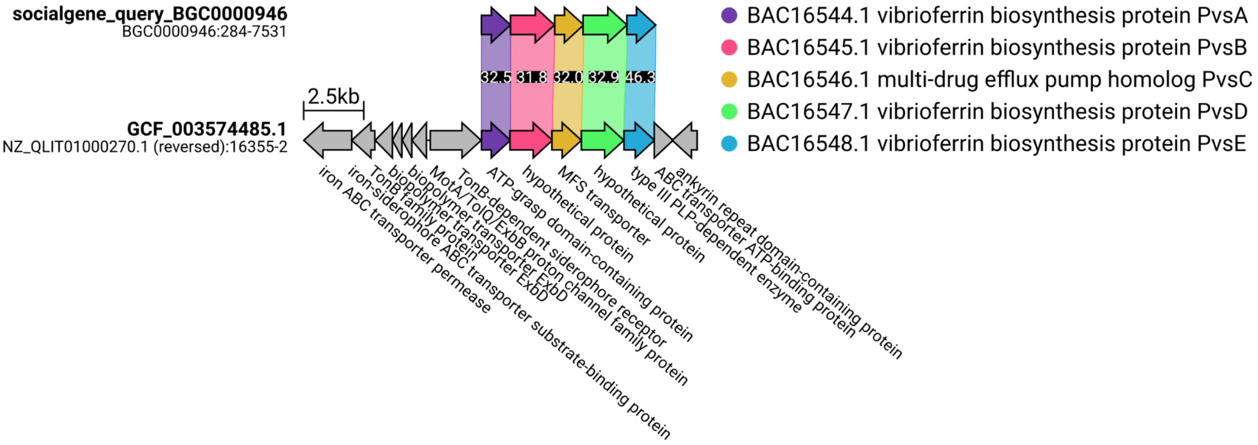


When searching for homologous gene clusters to BGC0000946 (vibrioferrin), the lowest median RBH identity of 32.5% belonged to *Facilibium subflavum* (GCF_003574485.1; NZ_QLIT01000270.1). The legend proteins and colors are derived from BGC0000946. The surrounding gene neighborhood suggests that the BGC is involved in metal homeostasis.

### Supplementary Figure 12: Putative vibrioferrin BGCs located on putative plasmids

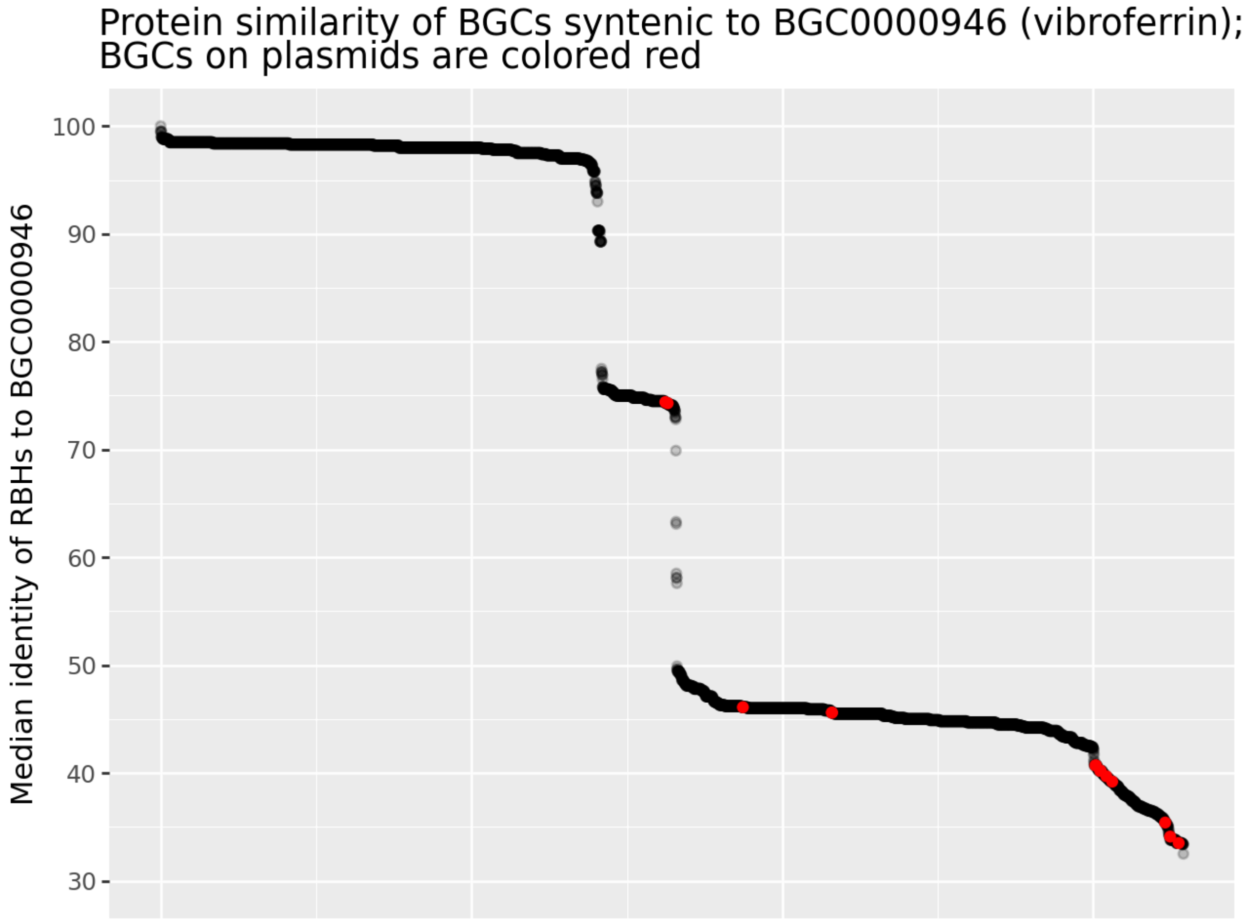


We were curious if there would be a tendency for BGCs near the cliffs to be seen on plasmids which might signal that a genetic transfer event occurred. This figure is identical to Fig. 4 except that for each BGC the nucleotide sequence identifier was used to query the SocialGene database and return results where the sequence was annotated in RefSeq as occurring on a plasmid, these are highlighted as red points. Though further study is needed, it is interesting that most BGCs occurring on plasmids are found after the first and second cliffs. Further studies could use SocialGene to find and cluster transposes flanking the BGCs using the MMSeqs2 clusters.

#### Supplementary Figure 13: Connecting genomes to BGCs with MMseqs2

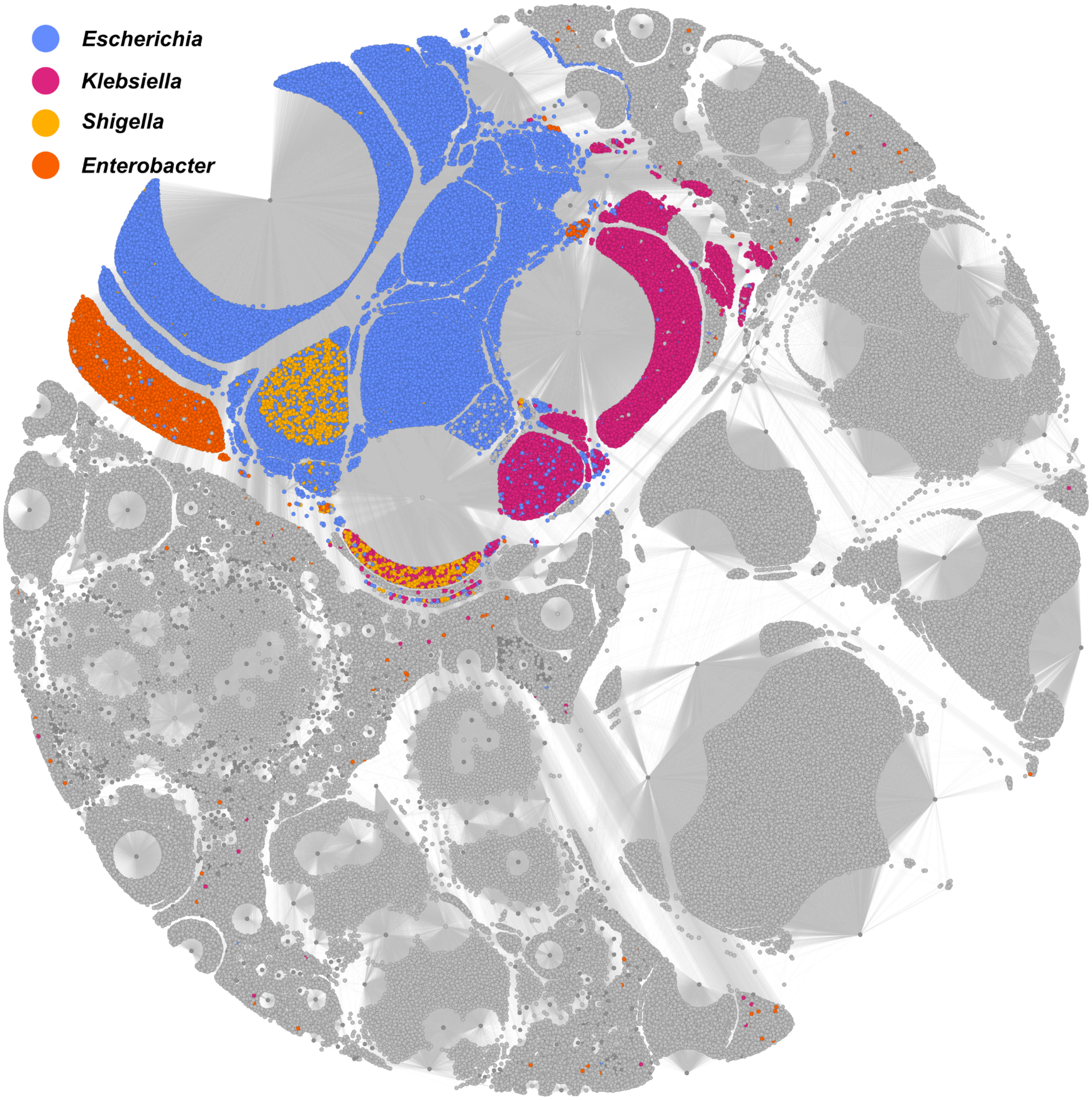


The full subgraph mentioned in Fig 5. Using a single Neo4j Cypher statement, new links (edges between nodes) were created between MIBiG BGCs (nodes) and genomes assemblies (nodes). New links were created where an assembly contained an antiSMASH predicted BGC whose proteins were at least 70% similar to at least 70% of a MIBiG BGC. The new subgraph consists of 366,404 relationships between 1,721 MIBiG BGCs and 158,479 genome assemblies.

#### Supplementary Figure 14: Connecting genomes to BGCs with MMseqs2 and highlighting culture collection organisms

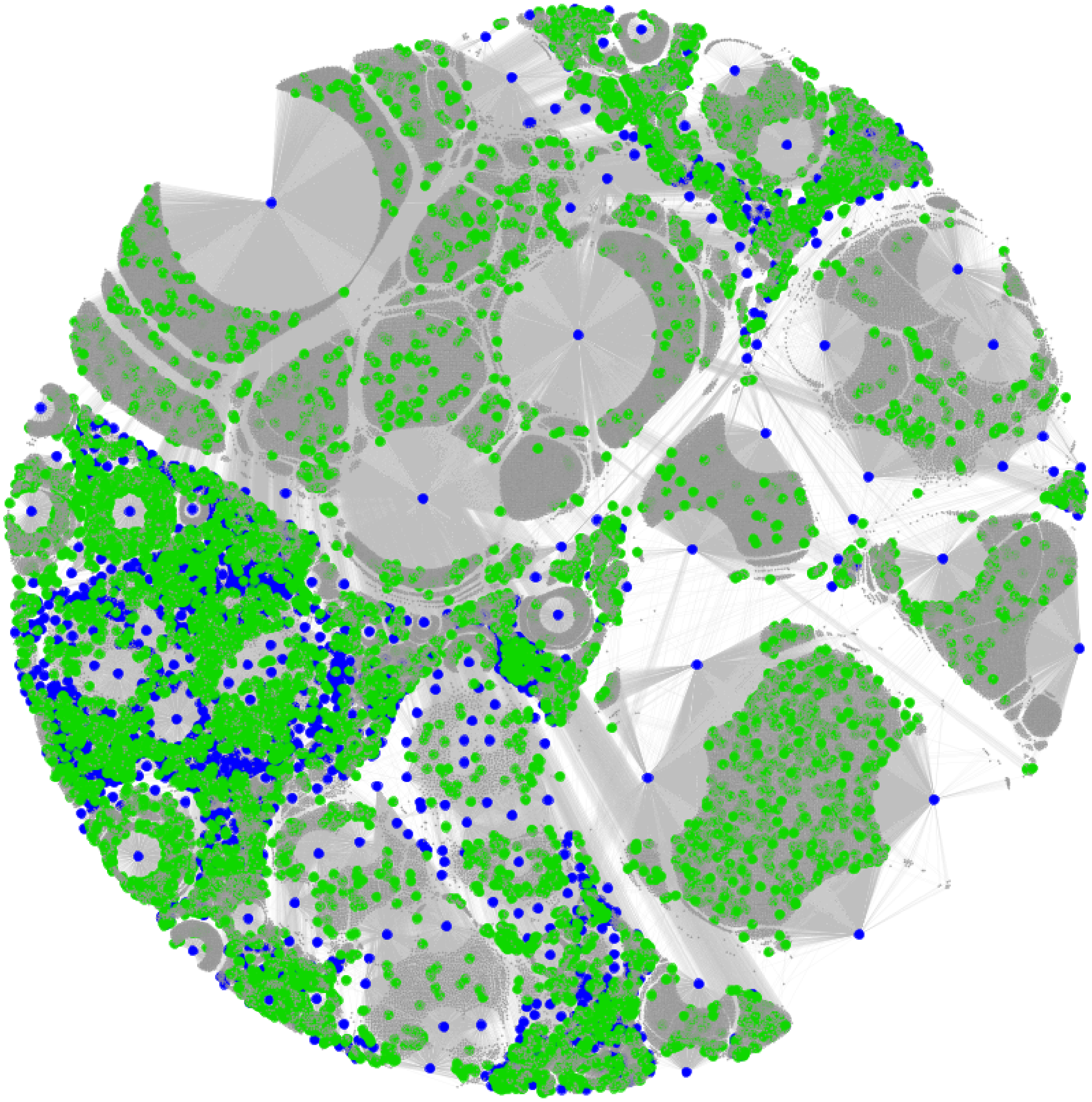

The same nodes and relationships as in Supplementary Fig. 13. The green nodes represent genome assemblies of strains that could be found in a strain collection (e.g. NRRL, ATCC, DSMZ, etc.); blue nodes represent MIBiG BGCs; gray nodes represent other genome assemblies. Note that in small scale (in this document) it isn't possible to faithfully represent the 160,200 nodes of the graph (i.e. it is overplotted). However, especially when displayed interactively, such analyses allow researchers to quickly identify strains that are available for testing their hypotheses.

### Supplementary Figure 15: Connecting genomes to BGCs with MMseqs2; displaying chemistry in Cytoscape

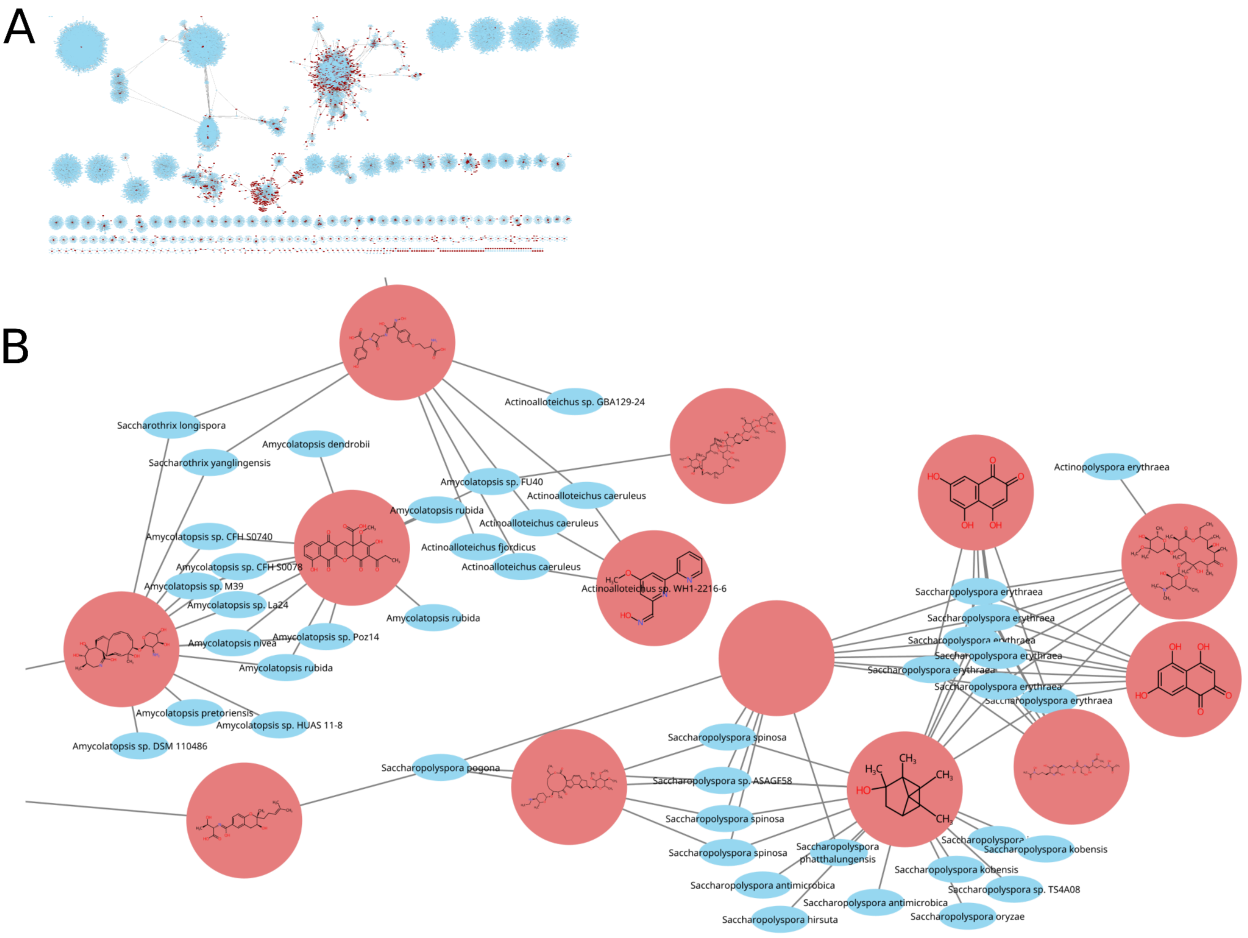


The same nodes and relationships as in Supplementary Figs. 13,14, but imported into Cytoscape directly from the database, using the Cytoscape Neo4j plugin. All MIBiG BGCs were connected to any RefSeq genome assembly which contained at least one antiSMASH 7 annotated region with proteins that matched at least 70% of the MIBiG BGC at 70% or greater identity. Protein similarity (identity) was determined by traversing MMseqs2 cluster relationships within the database (MMSEQS_90 and MMSEQS_70 edges). Panel A is a partial view of the resulting RefSeq assemblies and MIBiG BGCs (160,200 nodes) and the newly formed connecting edges. Not all nodes and relationships are displayed, due to limits of Cytoscape. Panel B is a random, zoomed-in subset of panel A, with genomes labeled with the organism's taxonomic names and MIBiG BGCs labeled by their associated chemical structures, where available, using the chemviz2 Cytoscape plugin.

### Supplementary Figure 16: Taxonomic placement of all MIBiG BGCs

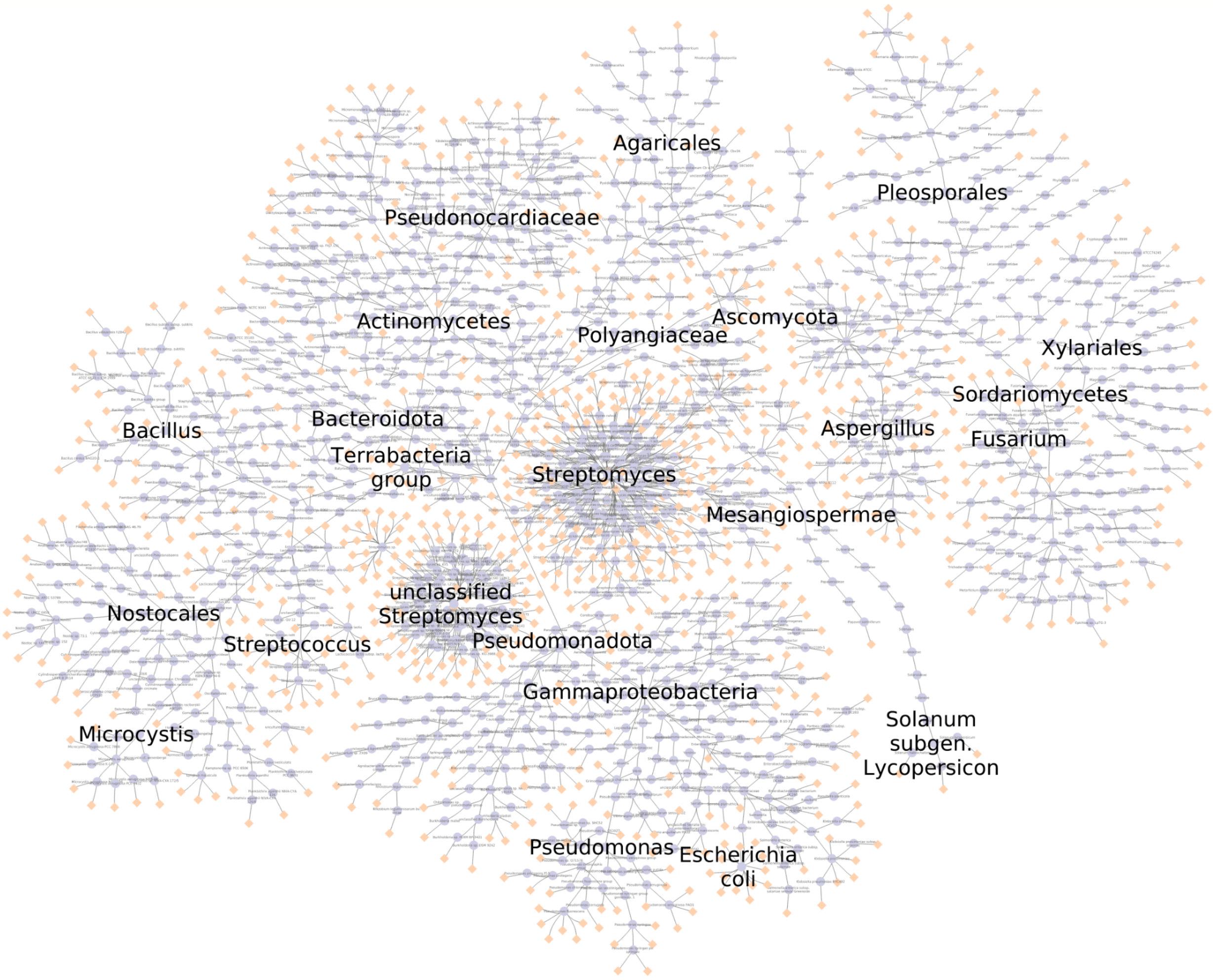


SocialGene's Nextflow workflow allows the incorporation of NCBI taxonomy into the resulting graph database. Here we show the taxonomic placement of all MIBiG BGCs within a SocialGene graph database. Blue nodes (circles) represent individual taxa from NCBI taxonomy and terminal yellow nodes represent MIBiG BGCs. Relationships (lines) between blue nodes represent taxonomic hierarchy (e.g. species, genus, family, etc.). Choice taxa were labeled in order to orient the reader. Combining analyses like this with Cytoscape chemistry plugins allows users to visualize chemical distribution across taxonomy, as does in-database chemical similarity and other cheminformatics.

### Supplementary Figure 17: MIBiG protein and pHMM graph

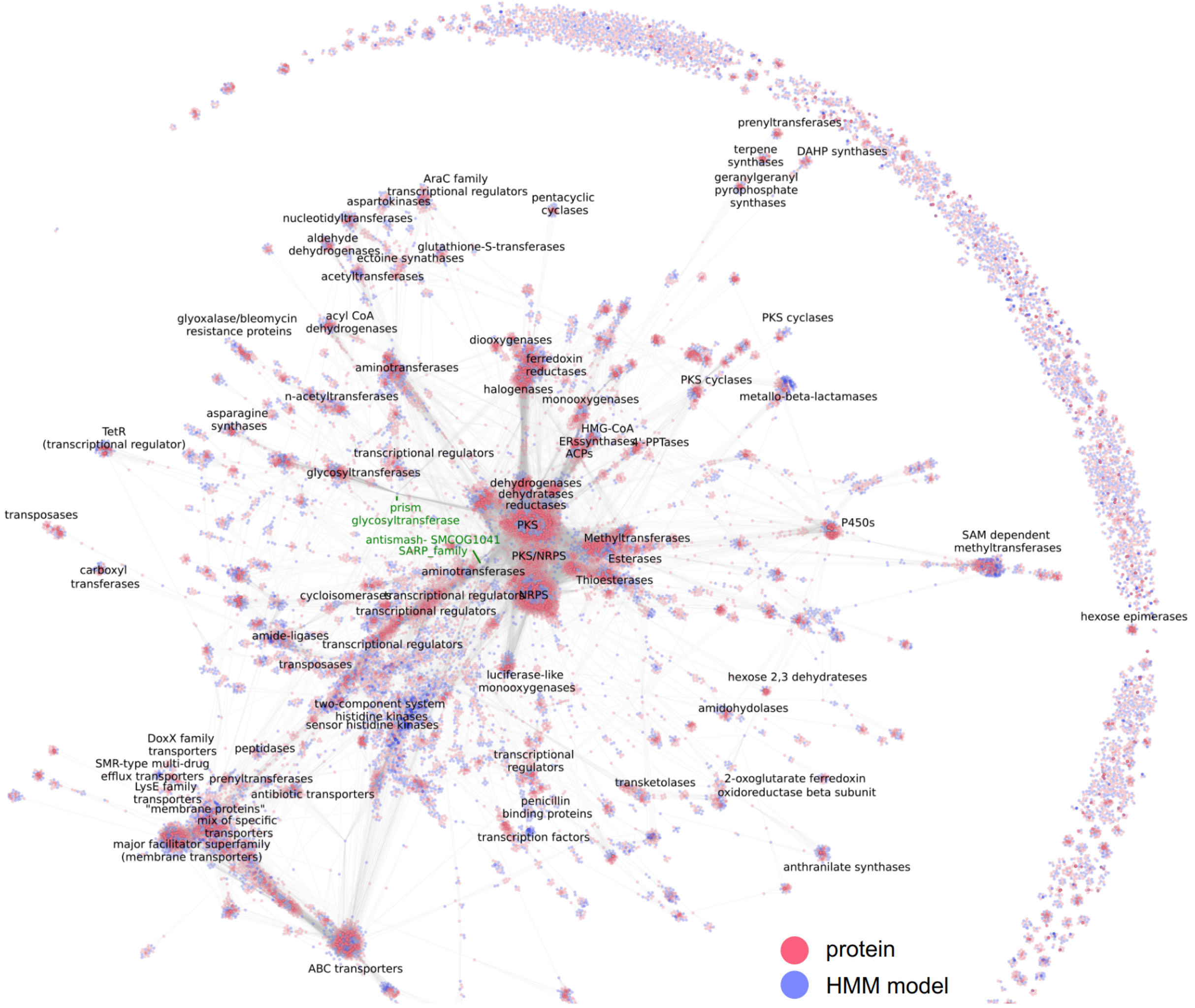


A minimal SocialGene database was created containing all MIBIG BGCs. Using Neo4j Cypher's "apoc.export.graphml.query" function proteins, pHMM models, and the links connecting the two were exported to a graphml. The graphml file was then imported into Gephi and arranged with ForceAtlas2[8]. Clusters of proteins and pHMM models were manually evaluated and found to be largely homogenous in function (based on protein names/descriptions). A subset of clusters were labeled according to their consensus protein name or class.
An interactive version of the graph is available at https://socialgene.github.io/mibigmap.

### Supplementary Figure 18: Targeted drug discovery cypher query

```
:auto
MATCH z1=(n:pfam {name:"Trp_halogenase"})←[:SOURCE_DB]-(h1:hmm),
      z2=(h1)-[:ANNOTATES]-(:protein)←[e1:ENCODES]-(n1:nucleotide)
WHERE n1.external_id STARTS WITH "BGC"
CALL {
    WITH n1, e1
    MATCH z3=(an1:hmm_source:antismash)←[:SOURCE_DB]-(:hmm)-[:ANNOTATES]→(p1:protein)←[e2:ENCODES]-(n1)
    MATCH z4=(an2:hmm_source:antismash)←[:SOURCE_DB]-(:hmm)-[:ANNOTATES]→(p1)
        WHERE an1.name ="Condensation"
            AND an2.name IN ["AMP-binding", "A-OX"]
            AND abs(e1.start - e2.start) < 10000
            AND e1.strand = e2.strand
    MATCH z5=(:hmm_source:amrfinder)←[:SOURCE_DB]-(:hmm)-[:ANNOTATES]→(p2:protein)←[e3:ENCODES]-(n1)
    WHERE abs(e1.start - e3.start) < 50000
            AND e1.strand = e3.strand
    RETURN z3, z4, z5
} in transactions of 1 rows
RETURN z1, z2, z3, z4, z5
```

This Neo4j Cypher query emulates the antiSMASH NRPS and halogenase BGC detection rules as copied below, with differing restrictions on distance between genes and the additional requirements of having a nearby antibiotic resistance gene and all the genes occurring on the same strand of DNA.

The following antiSMASH rules were copied from:
https://github.com/antismash/antismash/blob/fa46d2c822b4bcca99a6c13bb1bf38a844bd49d7/antismash/detection/hmm_detection/cluster_rules/strict.txt

RULE halogenated
    CATEGORY other
    DESCRIPTION Halogenases are frequently involved in secondary metabolite biosynthesis
    SUPERIORS polyhalogenated-pyrrole
    CUTOFF 5
    NEIGHBOURHOOD 10
    CONDITIONS Trp_halogenase

RULE NRPS
    CATEGORY NRPS
    DESCRIPTION non-ribosomal peptide synthase
    CUTOFF 20
    NEIGHBOURHOOD 20
    CONDITIONS cds(Condensation and (AMP-binding or A-OX))

### Supplementary Figure 19: Targeted drug discovery, halogenated NRP antibiotics

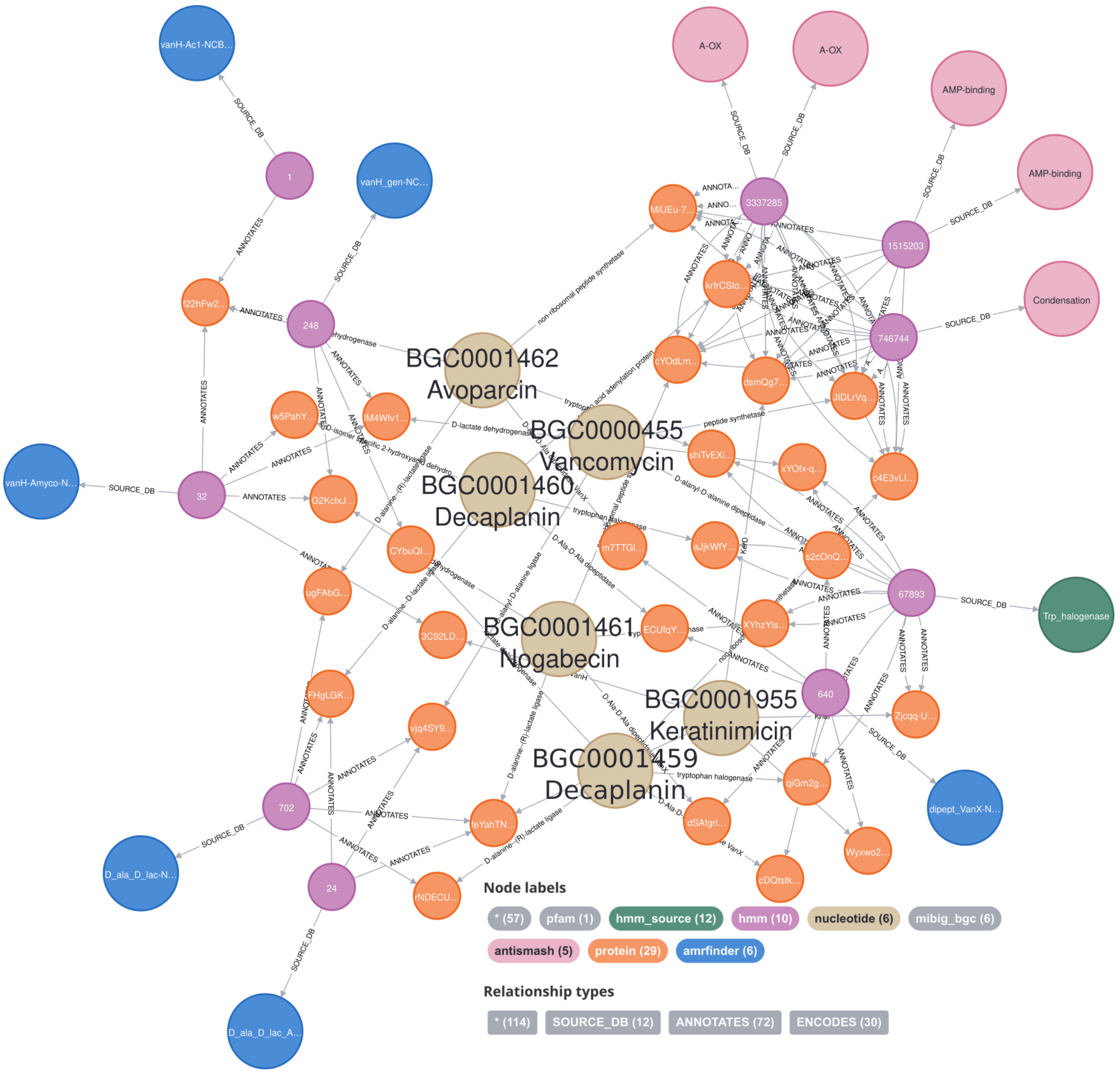


A screenshot of the Neo4j internet browser view of the results of the Cypher query in Supplementary Fig 18. The query successfully recovered halogenated NRP antibiotics from MIBiG (brown, labeled nodes).

### Supplementary Figure 20: SocialGene search for CAP superfamily protein containing BGCs

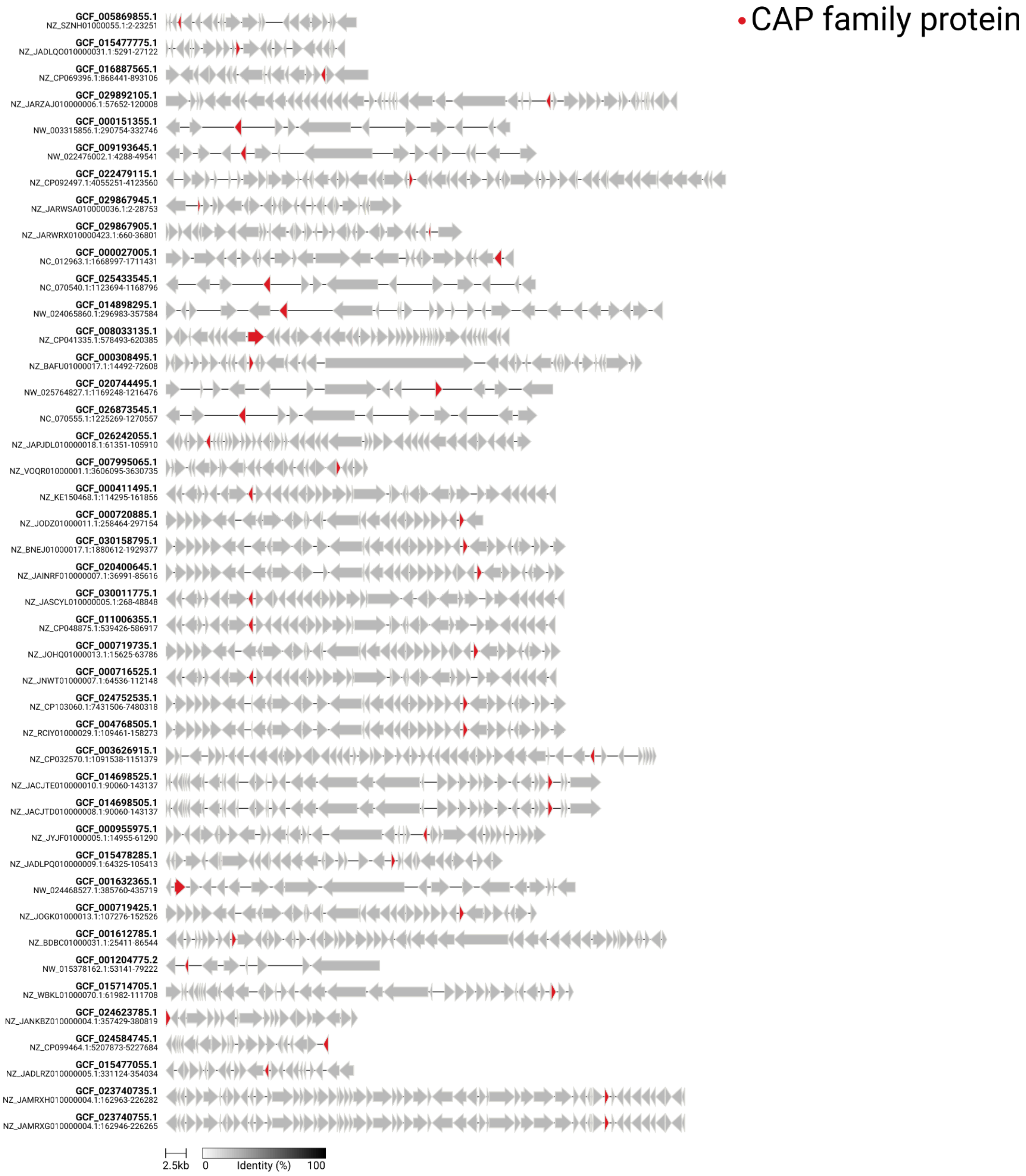


As proof of concept, >340,000 genomes (SocialGene RefSeq database) were searched for CAP protein containing BGCs (as annotated by antiSMASH 7). Utilizing SocialGene's Python library, the resulting putative BGC regions were converted to a clustermap.js plot and the putative CAP proteins highlighted in red.

#### Supplementary Figure 21: Screenshot showing the incorporation of NP Atlas into a SocialGene database

```
● (sgpy) chase@titan:~/Downloads$ sg_import_npatlas --input NPAtlas_download.json
  33372 Processing NPAtlas entries... ━━━━━━━━━━━━━━━━━━━━━━━━━━━━━━━━━━━━━━━━ 100% 0:02:30
  2024-03-08 08:35:13 INFO     Creating/Merging npatlas nodes in neo4
                      INFO     Connected to Neo4j database at bolt://localhost:7687
  2024-03-08 08:35:16 INFO     Created 33372 (:npatlas) nodes, set 467208 properties
  2024-03-08 08:35:18 INFO     Creating/Merging nodes linked to npatlas entries in neo4j
  2024-03-08 08:35:19 INFO     Created 13058 (:publication) nodes, set 78348 properties
                      INFO     Created 8 (:taxid) nodes, set 8 properties
  2024-03-08 08:35:20 INFO     Created 27196 (:gnps_library_spectrum) nodes, set 27196 properties
  2024-03-08 08:35:21 INFO     Created 96 (:assembly:mibig) nodes, set 96 properties
                      INFO     Created 1632 (:classyfire) nodes, set 1632 properties
                      INFO     Created 509 (:npclassifier_class) nodes, set 509 properties
                      INFO     Created 7 (:npclassifier_pathway) nodes, set 7 properties
                      INFO     Created 76 (:npclassifier_superclass) nodes, set 76 properties
  2024-03-08 08:35:28 INFO     Created 33339 (:chemical_compound) nodes, set 1366899 properties
                      INFO     Created 911 (:chebi) nodes, set 1822 properties
                      INFO     Linking npatlas entries and related nodes in neo4j
  2024-03-08 08:35:31 INFO     33372 relationships created (:npatlas)-[:HAS]->(:publication)
  2024-03-08 08:35:33 INFO     31481 relationships created (:taxid)-[:PRODUCES]->(:npatlas)
  2024-03-08 08:35:36 INFO     31887 relationships created (:npatlas)-[:HAS]->(:gnps_library_spectrum)
                      INFO     2511 relationships created (:assembly:mibig)-[:PRODUCES]->(:npatlas)
  2024-03-08 08:35:38 INFO     32888 relationships created (:npatlas)-[:LOWEST_CLASS]->(:classyfire)
  2024-03-08 08:35:40 INFO     32888 relationships created (:npatlas)-[:DIRECT_PARENT]->(:classyfire)
  2024-03-08 08:35:41 INFO     10694 relationships created (:npatlas)-[:INTERMEDIATE_NODES]->(:classyfire)
  2024-03-08 08:36:15 INFO     444892 relationships created (:npatlas)-[:ALTERNATIVE_PARENTS]->(:classyfire)
  2024-03-08 08:36:18 INFO     31400 relationships created (:npatlas)-[:IS_A]->(:npclassifier_class)
  2024-03-08 08:36:21 INFO     34822 relationships created (:npatlas)-[:IS_A]->(:npclassifier_pathway)
  2024-03-08 08:36:23 INFO     28759 relationships created (:npatlas)-[:IS_A]->(:npclassifier_superclass)
  2024-03-08 08:36:26 INFO     33371 relationships created (:npatlas)-[:IS_A]->(:chemical_compound)
  2024-03-08 08:37:37 INFO     801979 relationships created (:npatlas)-[:IS_A]->(:chebi)
```

SocialGene Python library's command line programs offer feedback and progress updates. Here NP Atlas is downloaded, parsed, and incorporated into a running SocialGene database.

#### Supplementary Figure 22: NP Atlas compounds linked by chemical similarity within a SocialGene database

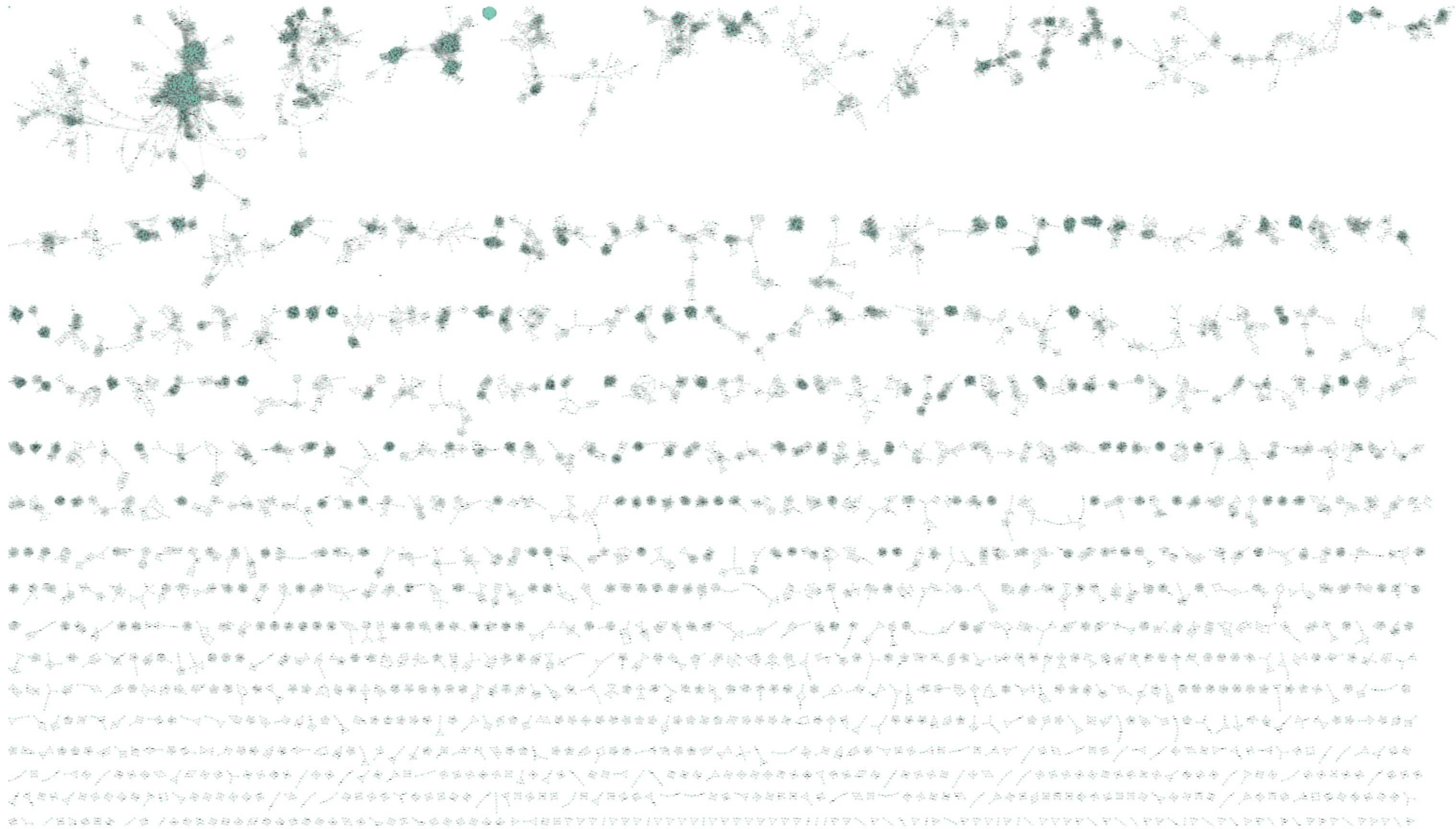

All NP Atlas compounds and all-vs-all chemical similarity relationships (Tanimoto) were imported into Cytoscape from a SocialGene database using the Neo4j Cytoscape plugin. As the number of genomes encompassing the taxonomic diversity of NP Atlas increases we expect more links and predictions can be made connecting BGCs to isolated compounds, and vice-versa. A zoom in is shown in Supplementary Figure 23.

### Supplementary Figure 23: Screenshot showing Tanimoto-similarity links between chemical compounds in a SocialGene database

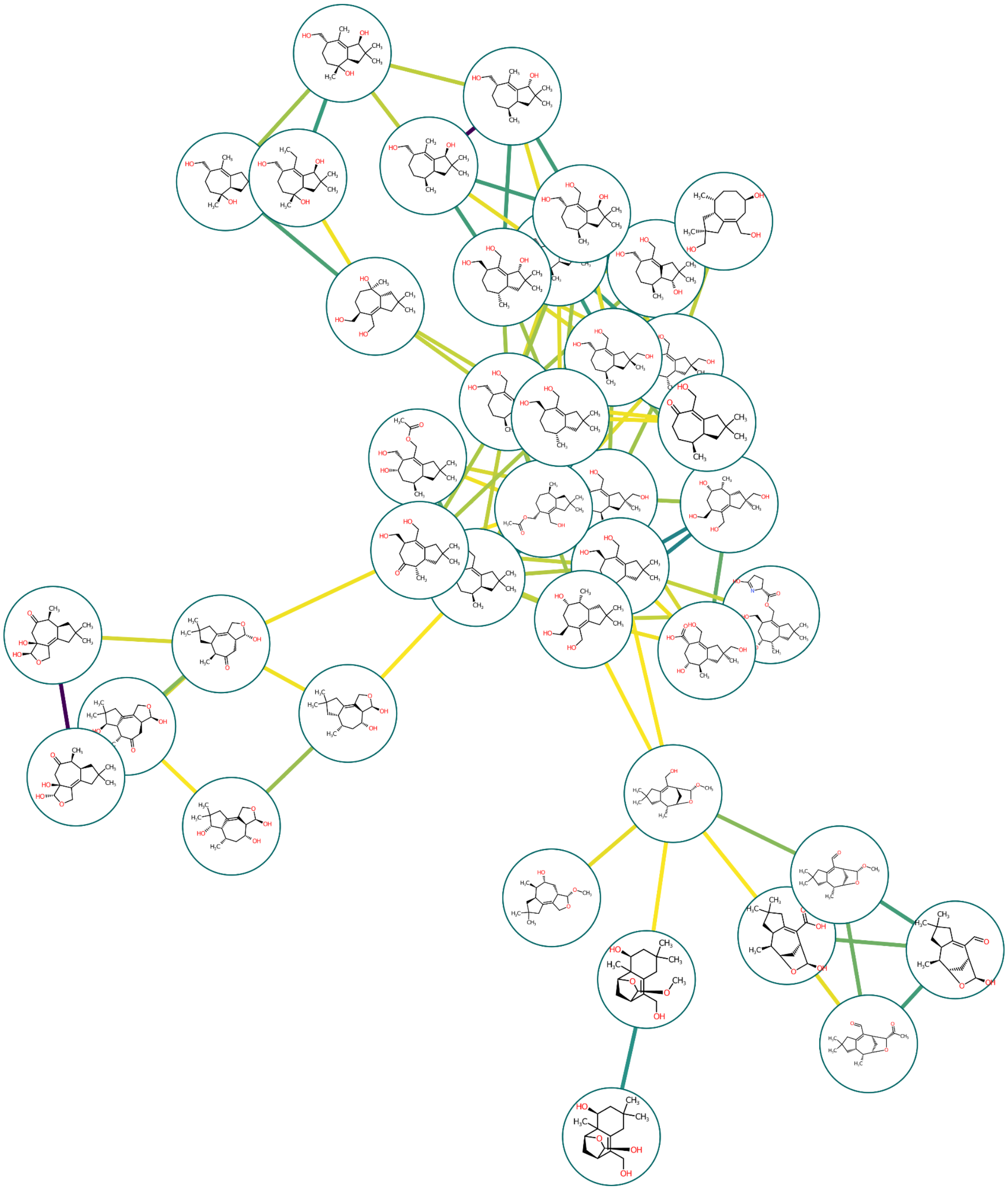

SocialGene has a command line function that calculates all-vs-all chemical similarity between all non-redundant chemical compounds in a running database. This allows users to create a coherent graph even when incorporating chemical structures from multiple databases and sources. The figure was generated using the Neo4j and chemviz2 plugins in Cytoscape and is a subset of Supplementary Figure 22.

Supplementary Figure 24: Screenshot showing the incorporation GNPS molecular networking results into a SocialGene database

| Metabolomics project identifier | Principal investigator | Submitter(s) | Nr of (meta)genomes | Nr of proteomes | Nr of growth conditions | Nr of extraction methods | Nr of instrumentation methods | Nr of links between genome and metabolome samples | Nr of links between biosynthetic gene clusters and MS/MS spectra |
|---|---|---|---|---|---|---|---|---|---|
| MSV000084723 | Cameron Currie & Tim Bugni | Marc G Chevrette | 120 | 0 | 1 | 1 | 1 | 122 | 0 |

```
(sgpy) chase@titan:~/Downloads$ sg_import_gnps --gnps_dirpath ProteoSAFe-METABOLOMICS-SNETS-V2-927974dc-view_all_clusters_withID_beta --map_path genome_to_file_map.csv
 2024-03-08 08:31:47 INFO     Connected to Neo4j database at bolt://localhost:7687                                  neo4j.py:62
 2024-03-08 08:31:50 INFO     Created 21930 (:ms2_spectrum) nodes, set 131580 properties                   neo4j_element.py:369
                     INFO     Created 63 (:gnps_library_spectrum) nodes, set 1633 properties               neo4j_element.py:369
                     INFO     Created 0 (:gnps_library_spectrum) nodes, set 0 properties                   neo4j_element.py:369
                     INFO     Created 1214 (:gnps_cluster) nodes, set 25695 properties                     neo4j_element.py:369
 2024-03-08 08:31:52 INFO     Created 122 (:mass_spectrum_file) nodes, set 366 properties                  neo4j_element.py:369
 2024-03-08 08:31:55 INFO     20027 relationships created (:ms2_spectrum)-[:CLUSTERS_TO]->(:gnps_cluster)  neo4j_element.py:538
 2024-03-08 08:31:59 INFO     21930 relationships created (:mass_spectrum_file)-[:HAS]->(:ms2_spectrum)    neo4j_element.py:538
                     INFO     Created 0 (:gnps_cluster) nodes, set 0 properties                            neo4j_element.py:369
                     INFO     Created 55 (:gnps_library_spectrum) nodes, set 55 properties                 neo4j_element.py:369
                     INFO     67 relationships created (:gnps_cluster)-[:LIBRARY_HIT]->(:gnps_library_spectrum) neo4j_element.py:538
                     INFO     Created 0 (:gnps_cluster) nodes, set 0 properties                            neo4j_element.py:369
                     INFO     Created 0 (:gnps_library_spectrum) nodes, set 0 properties                   neo4j_element.py:369
                     INFO     67 relationships created (:gnps_cluster)-[:LIBRARY_HIT]->(:gnps_library_spectrum) neo4j_element.py:538
 2024-03-08 08:32:00 INFO     Created 0 (:gnps_cluster) nodes, set 0 properties                            neo4j_element.py:369
                     INFO     Created 0 (:gnps_cluster) nodes, set 0 properties                            neo4j_element.py:369
                     INFO     2018 relationships created (:gnps_cluster)-[:MOLECULAR_NETWORK]->(:gnps_cluster) neo4j_element.py:538
                     INFO     Created 39 (:chemical_compound) nodes, set 1599 properties                   neo4j_element.py:369
                     INFO     46 relationships created (:gnps_library_spectrum)-[:IS_A]->(:chemical_compound) neo4j_element.py:538
                     INFO     Assemblies in GNPS results found in db: 84 of 84                                   parse.py:237
                     INFO     Assemblies in GNPS results not found in db: set()                                  parse.py:240
 2024-03-08 08:32:01 INFO     86 relationships created (:mass_spectrum_file)-[:ANALYSIS_OF]->(:assembly)   neo4j_element.py:538
                     INFO     GNPS molecular network has been integrated into the SocialGene Neo4j database        cli.py:146
```

#### Supplementary Figure 25: Connecting genomic and metabolomic data

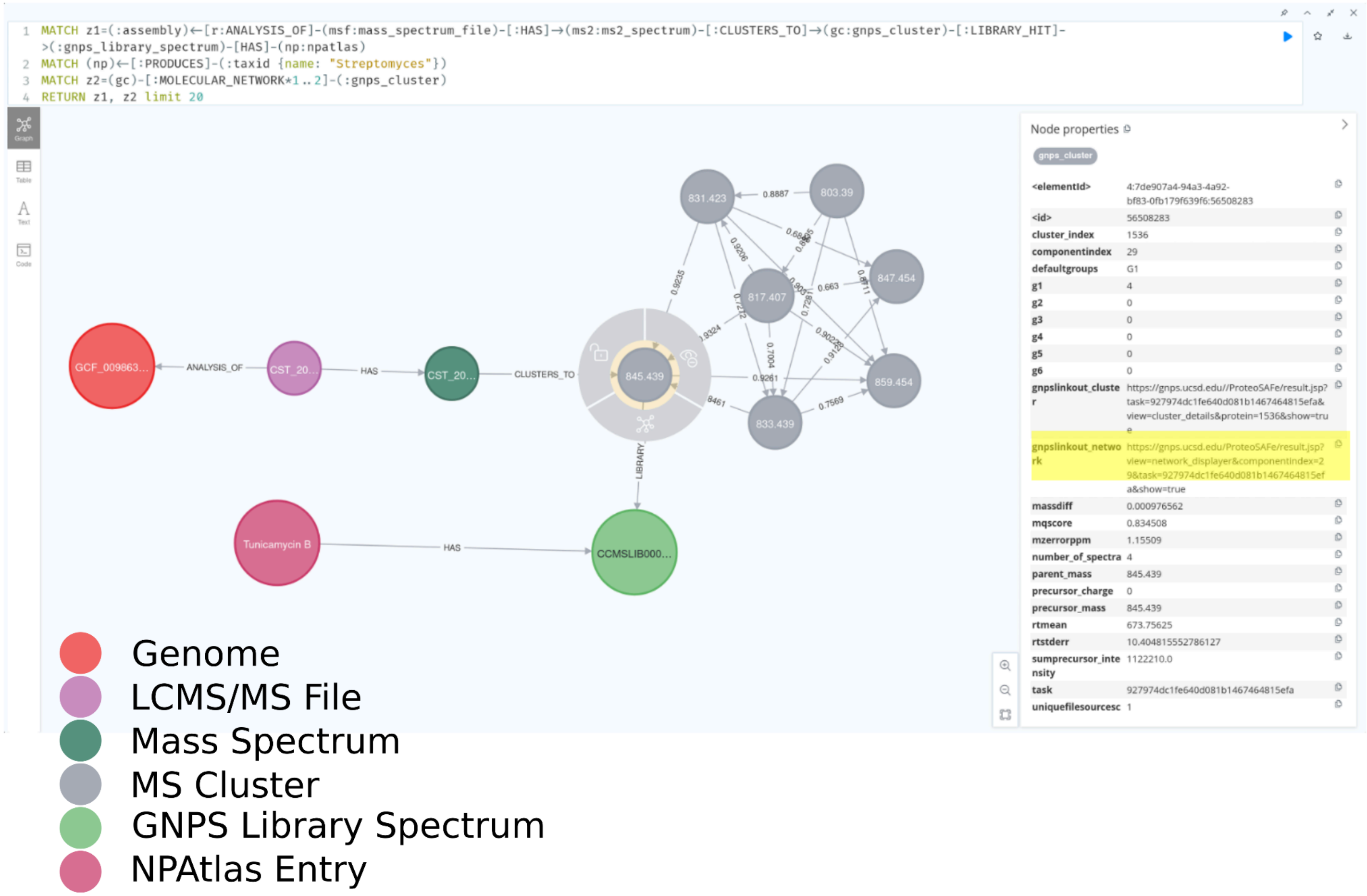


A subset of the resulting graph imported in Supplementary Fig. 24. The query identifies genomes associated with an MS cluster, that has a GNPS library hit, that is found in NP Atlas, where the NP Atlas entry was isolated from a Streptomycete. Not shown are additional links out from the genome assembly (e.g. nucleotide sequences, proteins, pHMM annotations, etc.) and NP Atlas (e.g. chemical ontology, taxonomic source, etc.).

### Supplementary Figure 26: SocialGene cross references MS clusters to GNPS website

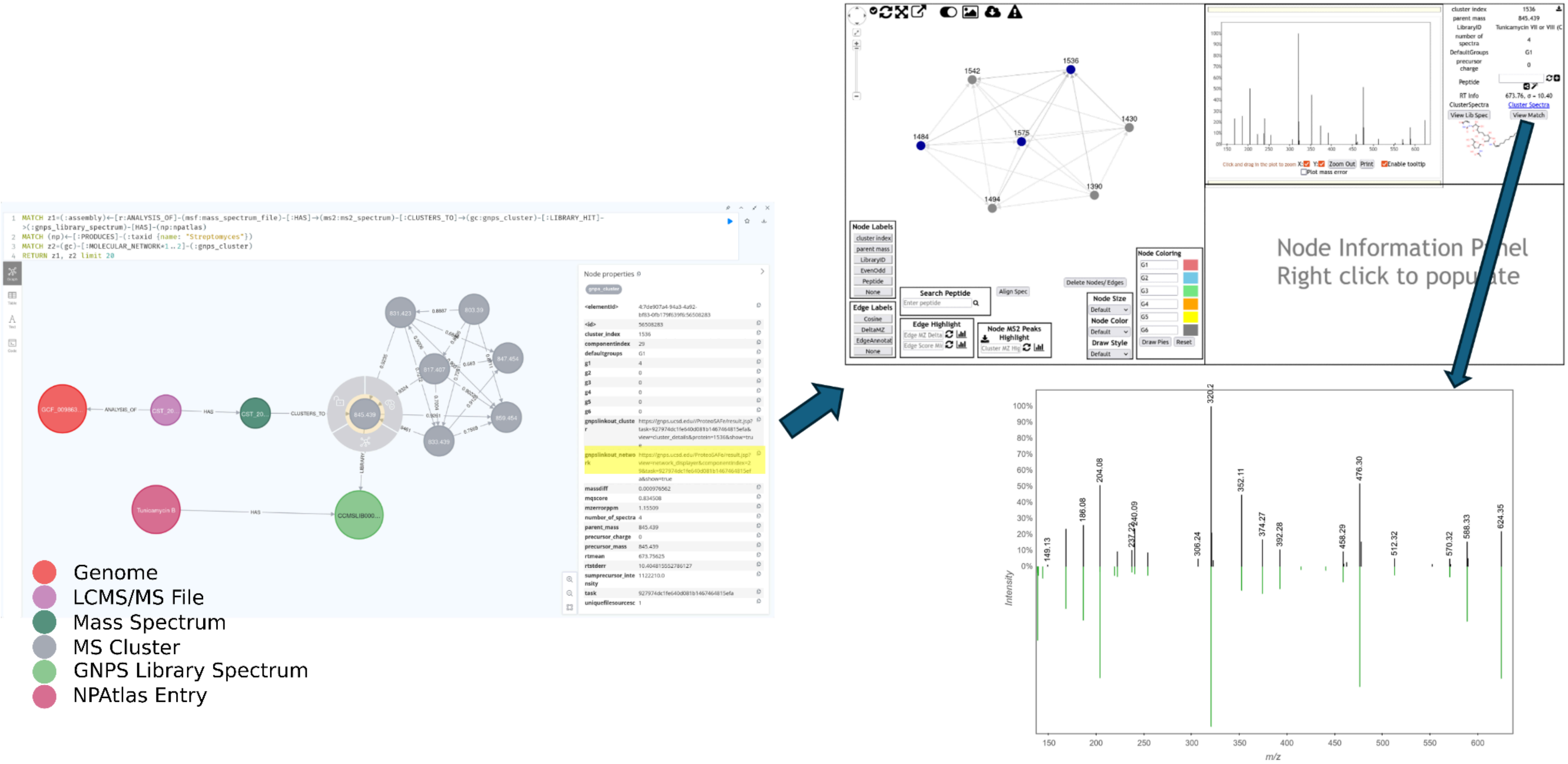


Here a mass spectrum cluster (gray node with additional ring), showing as a hit to a GNPS library spectrum, includes in its properties the url to view the cluster on the GNPS website where it was confirmed that the sample and library MS[2] spectra are nearly identical.

#### Supplementary Table 1: Types of antiSMASH-predicted BGCs containing a potential proteasome subunit, across >343,000 RefSeq genomes

| BGC Type | Count |
|---|---|
| "[ectoine]" | 257 |
| "[NRPS-like]" | 225 |
| "[T3PKS]" | 123 |
| "[terpene]" | 107 |
| "[T1PKS]" | 81 |
| "[NRPS, T1PKS]" | 74 |
| "[NRPS]" | 72 |
| "[arylpolyene, resorcinol]" | 69 |
| "[NRPS-like, T1PKS]" | 65 |
| "[arylpolyene]" | 57 |
| "[NRP-metallophore, NRPS]" | 38 |
| "[fungal-RiPP-like]" | 36 |
| "[T1PKS, hglE-KS]" | 34 |
| "[betalactone]" | 28 |
| "[RiPP-like]" | 27 |
| "[acyl_amino_acids]" | 19 |
| "[acyl_amino_acids, hserlactone]" | 17 |
| "[NRPS, NRPS-like, T1PKS]" | 16 |
| "[T2PKS]" | 15 |
| "[T1PKS, terpene]" | 15 |
| "[hglE-KS]" | 12 |
| "[redox-cofactor]" | 11 |
| "[RRE-containing]" | 11 |
| "[lanthipeptide-class-v]" | 11 |
| "[RiPP-like, terpene]" | 8 |
| "[hserlactone]" | 8 |
| "[LAP]" | 7 |
| "[NAGGN]" | 7 |
| "[thiopeptide]" | 7 |
| "[NRPS-like, T1PKS, transAT-PKS-like]" | 7 |
| "[T1PKS, T2PKS]" | 6 |
| "[NAPAA]" | 6 |
| "[NI-siderophore]" | 6 |

| | |
|---|---|
| "[T3PKS, terpene]" | 6 |
| "[CDPS]" | 5 |
| "[cyanobactin]" | 4 |
| "[proteusin]" | 4 |
| "[NRP-metallophore, NRPS, acyl_amino_acids]" | 4 |
| "[T1PKS, betalactone, hglE-KS]" | 4 |
| "[phosphonate]" | 4 |
| "[phosphonate-like]" | 4 |
| "[NRPS-like, betalactone]" | 3 |
| "[arylpolyene, resorcinol, terpene]" | 3 |
| "[NRPS, hserlactone]" | 3 |
| "[other]" | 3 |
| "[NRPS, betalactone, transAT-PKS]" | 3 |
| "[lassopeptide]" | 3 |
| "[NRPS, NRPS-like]" | 3 |
| "[thioamitides, thiopeptide]" | 2 |
| "[indole]" | 2 |
| "[NRPS, NRPS-like, PKS-like, T1PKS, transAT-PKS-like]" | 2 |
| "[PKS-like]" | 2 |
| "[NRP-metallophore, NRPS, T1PKS]" | 2 |
| "[NRPS, PKS-like, T1PKS, terpene]" | 2 |
| "[2dos]" | 2 |
| "[NI-siderophore, NRPS]" | 2 |
| "[cyclic-lactone-autoinducer]" | 2 |
| "[NRPS, T1PKS, terpene]" | 2 |
| "[LAP, proteusin, thiopeptide]" | 1 |
| "[NRPS-like, T1PKS, hglE-KS, transAT-PKS-like]" | 1 |
| "[thioamitides]" | 1 |
| "[NRPS, NRPS-like, T1PKS, transAT-PKS-like]" | 1 |
| "[NRPS-like, RiPP-like, T1PKS, transAT-PKS]" | 1 |
| "[epipeptide]" | 1 |
| "[NRPS-like, terpene]" | 1 |
| "[NRPS, NRPS-like, transAT-PKS]" | 1 |
| "[NRPS, T1PKS, fungal-RiPP-like]" | 1 |
| "[T1PKS, fungal-RiPP-like]" | 1 |
| "[NRP-metallophore, NRPS, NRPS-like, T1PKS]" | 1 |
| "[arylpolyene, ectoine, resorcinol]" | 1 |
| "[NRPS, NRPS-like, T1PKS, T3PKS, terpene]" | 1 |
| "[isocyanide]" | 1 |

| | |
|---|---|
| "[NRPS, NRPS-like, T1PKS, betalactone, transAT-PKS-like]" | 1 |
| "[NRPS-like, transAT-PKS]" | 1 |
| "[lanthipeptide-class-iv]" | 1 |
| "[T1PKS, hglE-KS, oligosaccharide]" | 1 |
| "[NRPS, fungal-RiPP-like, terpene]" | 1 |
| "[isocyanide-nrp]" | 1 |
| "[mycosporine-like]" | 1 |
| "[fungal-RiPP]" | 1 |
| "[NAPAA, NRPS-like]" | 1 |
| "[NRPS, T1PKS, T3PKS, terpene]" | 1 |
| "[LAP, NRPS, T1PKS, thioamitides, thiopeptide]" | 1 |
| "[LAP, thiopeptide]" | 1 |
| "[methanobactin]" | 1 |
| "[linaridin]" | 1 |
| "[NRPS, NRPS-like, T1PKS, phenazine]" | 1 |
| "[PKS-like, T1PKS, terpene]" | 1 |
| "[ranthipeptide]" | 1 |
| "[T1PKS, T2PKS, transAT-PKS]" | 1 |
| "[NRPS, fungal-RiPP-like]" | 1 |
| "[transAT-PKS-like]" | 1 |
| "[fungal-RiPP, fungal-RiPP-like]" | 1 |
| "[NAGGN, T1PKS]" | 1 |
| "[T1PKS, T3PKS]" | 1 |
